# CAM5, WRKY53, and TGA5 regulate defense gene expression mediated by the volatile organic compound ethyl vinyl ketone

**DOI:** 10.1101/2023.01.24.525215

**Authors:** Junqing Gong, Zhujuan Guo, Zhaoyuan Wang, Lijuan Yao, Chuanfang Zhao, Sheng Lin, Songling Ma, Yingbai Shen

## Abstract

Plants produce ethyl vinyl ketone (evk) in response to biotic stress, but the evk’s identification and downstream defense response remain unclear. In this paper, it is predicted by docking for the first time that evk can be recognized by RBOH protein and assist the electron transfer of RBOHD/RBOHF by binding to its FAD or NADPH binding site. Here, we show that evk treatment increased H_2_O_2_ and intracellular calcium concentrations in *Arabidopsis thaliana* mesophyll cells, as observed by confocal laser scanning microscopy and non-invasive micro-test technology, and that H_2_O_2_ signaling functioned upstream of Ca^2+^ signaling. Yeast two-hybrid, firefly luciferase complementation imaging, and *in vitro* pull-down assays demonstrated that the ACA8 (AUTOINHIBITED Ca^2+^-ATPASE, ISOFORM 8)–CML8 (CALMODULIN-LIKE 8) interaction promoted Ca^2+^ efflux to return Ca^2+^ levels to the resting state. Evk treatment led to the antagonism of salicylic acid (SA) and jasmonic acid (JA). CALMODULIN 5 (CAM5) positively regulates *WRKY53* expression, and CAM5 and WRKY53 positively regulate SA-related gene expression. These proteins physically interact and form a complex that is unlocked by Ca^2+^ to release WRKY53. An electrophoretic mobility shift assay and dual-luciferase reporter assay demonstrated that WRKY53 and TGA5 cooperate to enhance the expression of the defense gene *PATHOGENESIS-REALTED 1* (*PR1*) and that WRKY53 enhances the binding of TGA5 to the *PR1* promoter. This paper proposes a framework that evk, as a RES substance, can achieve plant’s ‘REScue’ through complete defense response.

## Introduction

The classical ligand receptor reversible perception mechanism usually includes the following steps: Biological production of specific hormones; hormones are released and transported to specific locations; hormones bind through transmembrane or soluble receptor proteins; it transmits perceived hormone signals through signal transduction process and mediates cell reaction; hormone cycle is used for the next round of signal perception and transmission, or hormone decomposition to prevent hormone signal transmission^1^. In the past decades, many previous studies have revealed how plants perceive and transduce JA to confer plant defense. It is known that JA is sensed by COI1 receptor and induces JAZ inhibitor degradation^2^, leading to rapid activation of JA response genes necessary for JA regulated plant response. Defensive response refers to a series of specific internal metabolic changes and external structural changes of plants to improve their survival ability against insect and pathogen invasion, including: external stimulus recognition, stimulus signal transduction, expression and regulation of defense genes, synthesis and accumulation of bioactive substances, defense effect realization and other biological processes. Evk is one of the plant volatiles. It is not clear how plants recognize evk to mediate the signal transduction and cause plant defense process. The event is still unknown.

Volatile organic compounds can be released from plants in response to insect feeding, mechanical injury, or infection. Plant volatiles often play major roles in regulating insect and microbe feeding on plants^3^. Volatiles containing α,β-unsaturated carbonyls are known as reactive electrophilic substances (RESs)^4,5^. Cells can benefit from lipid peroxidation and RES production while under stress. For instance, RESs modulate gene expression via a class II TGA transcription factor module and unknown signaling pathways. RES signaling promotes cell “REScue” by activating genes encoding detoxification enzymes, cell cycle regulators, and chaperones. Most biological activities of oxygenated lipid (oxylipin) in plants are mediated by either jasmonate (JA) perception or RES signaling networks^6^. Notably, numerous secondary compounds, including many phenolics and terpenes, exhibit electrophilic properties (such as α,β-unsaturated carbonyl groups), suggesting that such lipid-derived molecules may have been among the earliest direct defense chemicals to emerge^7^. Other secondary compounds exhibit behaviors that strongly suggest unique, defense-related functions, although their roles are unknown. One of these compounds is malondialdehyde (MDA), which has a variety of variants, each with unique characteristics^8^. While many other minor RESs are volatile and can be released after an attack, the protonated form of MDA is likely too reactive to escape from plants as a volatile. MDA enhanced endogenous JA and salicylic acid (SA) in Arabidopsis after *Botrytis cinerea* infection^9^. (E)-2-hexenal is an electrophilic compound that is synthesized and released quickly during bacterial infection or insect feeding^10,11,12^. Although methyl vinyl ketone is toxic to cells at high concentrations^13,14^, such substances might activate the expression of genes such as *HEAT SHOCK PROTEIN 101* (*HSP101*) in isoprene-producing plants. Furthermore, 12-oxo-phytodienoic acid (OPDA), an electrophilic chemical, can be employed as a signaling molecule to increase the expression of a wide range of genes other than JA-related genes^15,16,17^. Mattick and Hand were the first to discover ethyl vinyl ketone (evk) in soybean (*Glycine max* (L.) *Merr*.) homogenates^18^.

Evk is a small, volatile chemical molecule with an α, β-unsaturated carbonyl structure that activates the expression of defense genes in Arabidopsis (*Arabidopsis thaliana*). Evk was released as a pest-induced plant volatile when cabbage butterfly (*Pieris rapae*) was fed to Arabidopsis^19^. Soybean produced evk in a 13-HOPT-dependent manner when LIPOXYGENASE (LOX) activity was inhibited^20^. When damaged by freeze-thaw and oxygen stress, soybean leaves emit a substantial amount of evk^21^. The occurrence of floral end rot was reduced by increasing the evk content of tomatoes (*Solanum lycopersicum*) by treating the plants with selenium during the green blossom stage^22^. We previously demonstrated that evk-induced stomatal closure requires MULTIDRUG RESISTANCE-ASSOCIATED PROTEIN 4 (MRP4)-dependent extracellular ATP accumulation and subsequent H_2_O_2_ accumulation to regulate K^+^ efflux^23^. Evk is involved in many biological processes, but whether it activates defense genes (e.g., *PLANT DEFENSIN 1*.*2* [*PDF1*.*2*] and *PATHOGENESIS-RELATED 1* [*PR1*]) remains unknown.

Ca^2+^ acts as a crucial early signal during plant defense responses. When plants detect external signs of pathogen attack, such as oral elicitors from insects or bacterial flagella, Ca^2+^ from intracellular calcium stores (in the vacuole, Golgi apparatus, or endoplasmic reticulum [ER]) or extracellular calcium stores (in the interstitial spaces between the cell wall and plasmids) immediately flow into the cytoplasm^24,25^. The Ca^2+^/proton antiporter system and Ca^2+^-ATPases are responsible for returning Ca^2+^ to the calcium reserves once it has fulfilled its function as a second messenger^26^.

Two mechanisms in plants require both Ca^2+^ and H_2_O_2_: Ca^2+^-induced ROS generation and ROS-induced Ca^2+^ release. Co-expressing *AtSRC2* (*SOYBEAN GENE REGULATED BY COLD 2*) and *RBOHF* in human embryonic kidney (HEK293) cells enhanced the production of ROS mediated by a Ca^2+^-dependent ROS burst^27^. Under salt stress, Ca^2+^ and ROS (whose production relies on AtRBOHD) concurrently undergo long-distance transmission^28^. When Ca^2+^ binds to the NADPH oxidase EF-hand domain, the chiral structure of the EF-hand undergoes a conformational change that is required for RBOHD activation. ROS directly activate or inhibit the activity of Ca^2+^ channels or pumps that control Ca^2+^ concentrations, representing the primary source of ROS-induced Ca^2+^ release^29^.

The w-box sequence is frequently present in the upstream regulatory regions of genes activated by SA, as well as genes related to plant disease resistance, the damage response, and senescence. Various biological and abiotic stress reactions are tightly associated with WRKY transcription factors^30-35^. WRKY53 positively regulates resistance to *Pseudomonas syringae*^36^. WRKY53 and EPITHIOSPECIFIER PROTEIN (ESR) mediate the antagonistic relationship between pathogen resistance and aging^37^, which likely involves the JA/SA balance. EMSAs (electrophoretic mobility shift assays) demonstrated that the DNA binding ability of WRKY53 is inhibited by ESR in the presence of WRKY53 *in vitro*^38,39^. WRKY53 positively regulates stomatal opening and boosts starch metabolism by lowering the amount of H_2_O_2_ in guard cells and directly binding to the *QUA-QUINE STARCH* (*QQS*) promoter^40^. In Arabidopsis, the presence of w-boxes in the *ALLENE OXIDE SYNTHASE* (*AOS*) promoter is required for binding by JASMONATE-ASSOCIATED VQ MOTIF GENE 1 (JAV1) and WRKY51^41^.

Calmodulin (CaM) responds to a range of stress and plant hormonal cues, including H_2_O_2_ and SA, by selectively binding to the CGCG-box sequences in the promoters of its target genes and activating their transcription^42,43^. Ca^2+^/CaM-mediated regulatory signals have long been known to play essential roles in maintaining H_2_O_2_ homeostasis, plant growth and development, plant-microbial symbiosis, immunological response, and freezing tolerance. A fascinating question about the underlying cell signal transduction network is how plants begin with the release of the simple second messenger intracellular Ca^2+^ ions in response to a variety of external stimuli, form Ca^2+^/CaM signals by binding to CaM, and interpret these signals into various physiological response processes accurately and efficiently^44,45^.

Special electrophile structures can also covalently modify proteins, including cyclopentenone, an important regulator of plant development and responses to biotic and abiotic stress^46,47^. Its α, β-unsaturated carbonyl structure confers chemical reactivity to mercaptan, which is essential for protein activity^4,6,48^. TGACG MOTIF-BINDING FACTOR 2 (TGA2) enhances cyclopentenone-induced changes in gene expression. TGA2 activity is regulated in part by RES-dependent post-translational modification. TGA2 is covalently modified by Prostaglandin A1 (PGA1)-biotin (a RES not found in plants) and OPDA (an important plant RES). Extreme temperature and pH conditions can make covalently modified TGA2 vulnerable^49^. The carbonyl cyclopentenone ring in PGA1 may react with protein thiol (Cys) and/or primary amine (Lys, Arg)^50^. The primary production of conjugates relies on mercaptan modification, which occurs when PGA1-biotin modifies TGA2 at Cys-186 by Michael addition. A deletion mutant of a type II TGA transcription factor (*tga2, tga5*, or *tga6*) is unable to trigger PPA1 and OPDA production. When TGA2,5,6 mutations occur, the up regulation of α,β-unsaturated carbonyl structure-mediated gene decreases by 60%. A putative binding site (TGACG) for basic leucine zipper (bZIP) transcription factors of the TGA family with a TGA motif is present in the promoters of approximately 50% of cyclopentenone-induced genes^50^. These findings suggest that TGA transcription factors might function in response to RES^51^.

TGA-binding sites are found in the promoters of several genes and can react to a variety of signals during responses to pathogens, trauma, and xenobiotic stress^52-54^. Most JA-induced genes are not TGA dependent, while the majority of TGA-dependent detoxifying genes are not induced by JA^55^. In various yeast two-hybrid screening experiments, NPR1 was shown to interact with the TGA subclass of bZIP transcription factors^56,57^. These TGA transcription factors can bind to the as-1 element in the *PR1* promoter, which is required for the response of this gene to SA. The TGA2-NPR1 complex is better able to form in the presence of SA. Active NPR1 increases the binding of TGAs to the as-1 sequence in the *PR1* promoter, thereby increasing its expression^58^.

TGA3 and WRKY53 physically interact and regulate the activity of the *Cestrum yellow leaf curling virus* (CmYLCV) promoter. The two distinct types of TGA and WRKY transcription factors are known to interact^59^, but the nature if this interaction is not well understood. To regulate the expression of *PR1* in response to SA induction, the transcription factors TGA2 and AtWRKY50 bind to as-1 and the w-box in the *PR1* promoter, respectively (at an interval of 8 bp)^60^. In the presence of NPR1, the transcription of the CmYLCV promoter is regulated by SA only in the presence of TGA3 and WRKY53. SA-mediated *PR1* transcription in Arabidopsis is significantly affected by the TGA/NPR1 interaction^61,62^.

Here, we show that evk can be recognized by RBOH protein and evk treatment increases intracellular H_2_O_2_ and Ca^2+^ concentrations in leaves, as revealed using non-invasive micro-test technology (NMT) and confocal laser scanning microscopy. Together, CML8 and ACA8 transport Ca^2+^ out of the cytoplasm and lower cytoplasmic Ca^2+^ concentrations. Additionally, RBOHD and RBOHF influence the ROS burst as well as the rise in intracellular Ca^2+^ concentrations and the outflow of calcium ions. Therefore, evk functions as a signal that causes variation in Ca^2+^ and H_2_O_2_ concentrations in Arabidopsis leaves. We performed EMSAs and yeast two-hybrid, firefly luciferase complementation imaging, *in vitro* pull-down, and luciferase activity assays to explore how CAM5 and WRKY53 govern *PR1* and *PDF1*.*2* expression. High Ca^2+^ concentrations caused the CAM5-WRKY53 complex to disassemble, which released WRKY53. The released WRKY53 can interact with TGA5 in the nucleus, thereby increasing its ability to associate with the *PR1* promoter. WRKY53 binds to the *PDF1*.*2* promoter via its w-box, but the released WRKY53 can prevent TGA5 from binding to as-1 in the *PDF1*.*2* promoter. We also performed a dual-luciferase reporter assay to explore how both WRKY53 and TGA5 increase the transcription of *PR1*. WRKY53 and TGA5 more effectively triggered the expression of *PR1* together than separately.

*PDF1*.*2* was downregulated by WRKY53 but upregulated by TGA5.

## Materials and methods

### Computational detials

Our experimental results indicated that the ethyl vinyl ketone (evk)-mediated ROS burst is dependent on RBOHD and RBOHF, but how evk binds with RBOHD/RBOHF is still unknown. To explore the mode of evk on RBOHD or RBOHF, their interactions were mimicked with Molecular Operating Environment (MOE) 2020.09 software (Chemical Computing Group Inc., Montreal, Canada).

### Homology modeling of RBOHD and RBOHF

The crystal structure of AtRBOHD and AtRBOHF had not been reported, therefore 3D structures of them were built with homology modeling firstly. Homology modeling of AtRBOHD and AtRBOHF were executed with SwissModel, which is a fully automated protein homology modeling server (https://swissmodel.expasy.org/). The sequences of AtRBOHD and AtRBOHF were downloaded from UniProt website (Uniprot code: Q9FIJ0 and O48538). The 3D structure of AtRBOHD was built, with the QMEANDisCo Global is 0.60, GMQE is 0.48, and 30.01% sequence identity with human DUOX1-DUOXA1. And the 3D structure of AtRBOHF was built with the QMEANDisCo Global is 0.59, GMQE is 0.47, and 28.38% sequence identity with human DUOX1-DUOXA1.

### Binding site exploration and docking simulations

The possible binding sites of AtRBOHD and AtRBOHF were explored with Site Finder module in the MOE software. The top two recommended binding sites were used as binding sites of AtRBOHD and AtRBOHF, corresponding to the FAD and NADPH binding site of the highly homologous human DUOX1-DUOXA1 (PDB code, 7D3F). Docking were carried out with default parameters (Placement setting as Triangle Matcher, Refinement setting as Rigid Receptor) to explore the binding interactions between evk and AtRBOHD and AtRBOHF.

### Plant materials and culture conditions

Wild-type (WT) Arabidopsis accession Col-0, the *cam5* mutant SALK_007371 (At2g27030), and the *wrky53* mutant SALK_034157 (At4g23810) were used as plant materials. The *wrky53* mutant was kindly provided by Prof. Diqiu Yu (Xishuangbanna Tropical Botanical Garden, Chinese Academy of Sciences, China). The *cml8-1* mutant SALK_022524C (At4g14640) and the *cml8-2* mutant SALK_114570C (At4g14640) were used as plant materials. Seeds were stratified at 4℃ for 2 days in the dark. After stratification, seeds were surface-sterilized for 4 min in 75% (v/v) ethanol, washed four times in sterile water, sown on autoclaved soil mixture, and placed in an incubator (Percival model: I-36vl). Plants were grown on soil at 21–23℃ and 70% relative humidity with a light intensity of 80–110 µmol m^−2^ s^−1^ under long-day (16 h light/8 h dark) conditions. The plants used for insect inoculation were grown for 4 weeks, and the seedlings used in all other experiments were grown for 2 weeks prior to treatment.

### Evk treatment

Arabidopsis plants were treated with evk (≥ 97%, purchased from Sigma) via fumigation in glass bell jars (height: 12.5 cm; diameter: 15 cm). Cotton balls 1 cm in diameter were soaked in evk dissolved in ethanol (evk at a final concentration of 5 µM) and hung in the bell jars; cotton balls soaked in the same volume of ethanol were used as control. After adding the cotton balls, the bell jars were immediately sealed with Vaseline petroleum jelly.

### Ca^2+^ flux measurements in mesophyll cells

Ca^2+^ flux in mesophyll cells was measured using the non-invasive micro-test technique (NMT, BIO-001A, Younger USA). Before the test, small cuts were made in the leaves of 3-week-old Arabidopsis plants. The leaves were soaked in test solution (0.1 mM KCl, 0.1 mM CaCl_2_, 0.1 mM MgCl_2_, 0.5 mM NaCl, 0.3 mM MES, 0.2 mM Na_2_SO_4_, pH 6.0) for approximately 30 min. Silanized glass micropipettes (2-to 4-µm aperture) were filled with electrolyte solution (100 mM CaCl_2_) and front-filled with a selective liquid ion exchange (LIX) cocktail to approximately 10 µm. The glass micropipettes were connected to the NMT system with a silver chloride wire, and electrodes with Nernstian slopes of 28 ± 5 mV log^−1^ were used.

The Ca^2+^ concentrations in mesophyll cells were measured near the cells and 30 µm away from the cells at 0.2 Hz. Each leaf was tested for 2 min before evk treatment (pre-evk treatment; pre). After evk was quickly added to the test solution to a final concentration of 10 µM, data were collected for approximately 2.5 min as the evk response peak (peak) group. Data were then collected from the post-evk response (post) group to indicate the end of the reaction. The final flux values are reported as the mean of eight individual plants per treatment. The Ca^2+^ flux was calculated as:

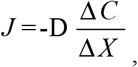

where *J* is the flux of Ca^2+^ (pmol cm^−2^ s^−1^), D is the diffusion coefficient (0.79 ×10^−5^ cm^−2^ s^−1^), Δ*C* is the difference between the concentrations near and far from the cells, and Δ*X* is 30 µm. Each group contained six to eight replicates.

### Ca^2+^ fluorescence measurements in mesophyll cells

Prior to treatment with Fluo3-AM (10 µM), Arabidopsis leaves were soaked in test solution (the same solution used for the Ca^2+^ flux experiment). The leaves were then soaked in Fluo3-AM solution to label Ca^2+^ ions for 1 h in the dark in an incubator. The leaves were washed and placed in fresh test solution. Ca^2+^ fluorescence was detected with a confocal laser scanning microscope (Leica TCS SP8). The excitation wavelength was 488 nm, and the emission wavelength was 530 nm. Before adding evk (0 min), the basal level of Ca^2+^ fluorescence in mesophyll cells was measured. Following the addition of evk, fluorescence was measured every 30 s for 150 s. Each group contained 20–40 cells.

The increase in Ca^2+^ fluorescence was calculated as (X – a) / a * 100%, where X is the fluorescence of evk-treated samples and a is fluorescence at 0 s. The value for each group was calculated separately.

### H_2_O_2_ fluorescence measurements in mesophyll cells

To observe transient evk responses, H2DCF-DA was used to measure changes in H_2_O_2_ levels in Arabidopsis mesophyll cells. Following incubation in the test solution, the leaves were incubated in 50 µM H2DCF-DA for 12 min at 25℃ in the dark. Confocal laser scanning microscopy (Leica TCS SP8, Leica Microsystems, Wetzlar, Germany) was used to detect H_2_O_2_ fluorescence with excitation at 490 nm and emission at 530 nm. H_2_O_2_ fluorescence measurements were obtained at 0 min (before the addition of evk) and every 3 min for 12 min. Each group contained 30 cells.

### RT-qPCR

For quantitative RT-PCR, total RNA was extracted from the samples using an RNA extraction kit, and first-strand cDNA was generated using a reverse transcriptase kit (Takara). qPCR was performed in a 7500 fast real-time PCR system (Applied Biosystems, Foster City, California, USA) using a Power SYBR Green PCR Master Mix kit (Applied Biosystems, Foster City, California, USA), and the 2^−ΔΔCt^ method was used to calculate relative gene expression levels. The expression levels of several important genes in the JA and SA were measured, See Supplementary 1 for primer information.

### Measuring JA/SA levels in leaves

JA and SA levels were measured in Arabidopsis leaves in six experimental groups (WT control group and WT evk group), and analysis was performed on an Agilent 1290 system with AB SCIEX-6500Qtrap. Each group had three biological replicates.

### Arabidopsis protoplast isolation and transformation

Arabidopsis protoplasts were isolated from leaf tissues of 14-day-old seedlings. Cut leaves into 0.5-1 mm thin strips, and soak them in 0.5 M mannitol solution for 30 min. Take out the sliver and put it into the enzymolysis solution (0.4 M mannitol, 20 mM KCl, 20 mM MES, 1.5% cellulase R-10, 0.4% macrozyme, 10 mM CaCl_2_) to avoid light, 23 ℃, 50 rpm, and enzymolysis on the shaking table for 3 hours. When the hydrolysate turns green, gently shake the culture dish to release protoplasts. Use a 40 µm cell sieve to filter the protoplasts into a centrifuge tube. Add an equal volume of W5 solution (2mM MES pH7.5, 154mM NaCl, 125mM CaCl_2,_ 5mM KCl). Centrifuge at 4 ℃ for 2 min at 100 × g. Add an equal volume of W5 solution. Repeat the washing once. Shake the centrifuged protoplasts gently and suspend them on ice for 30 min. Add an appropriate amount of precooled MMG solution (4 mM MES pH 5.7, 0.4 M mannitol and 15 mM MgCl_2_) to wash the protoplast, centrifuge for 1min, discard the supernatant, add an appropriate amount of MMG solution to suspend the protoplast, and transform PEG: the following steps are to take 10 µg of plasmid at 23 ℃ and put it into a 1.5ml centrifuge tube, add 100 µl of protoplast extract, and then add 110 µl of PEG conversion solution, mix it well, place it at 23 ℃ and incubate it in dark for 30 min, and transform it, the transfection mixture was diluted with 400 µl W5 solution to stop the transfection process followed by centrifugation at 100 × g for 1min at room temperature, incubation overnight in the dark in a 23 ℃ growth chamber.

### Dual-luciferase reporter assay

The *PR1* and *PDF1*.*2* promoters were cloned into pGreenII 0800-LUC and the *WRKY53, CAM5*, and *TGA5* coding sequences were cloned into pGreenII 62-SK. The resulting plasmids were transformed into Agrobacterium (*Agrobacterium tumefaciens*) strain GV3101 and used to infiltrate *Nicotiana benthamiana* leaves. Luciferase activity in infiltrated *N. benthamiana* leaves was imaged using a molecular imaging system (LB983). Firefly luciferase (LUC) and Renilla luciferase (REN) activities were measured using a Dual Luciferase Reporter Gene Assay Kit with a GloMax 96 Luminometer.

### Electrophoretic mobility shift assay (EMSA)

The w-boxes in *PR1, PDF1*.*2* promoter, and the as-1 element in the *PDF1*.*2* promoter were used to generate 3′-biotin-labeled probes. The CDS of CAM5 was cloned into pMAL-C2X with BamHI/SalI, the CDS of TGA5 was cloned into pMAL-C2X with BamHI/EcoRI, and the CDS of WRKY53 was cloned into MAL-C2X with BamHI/SalI. MBP, CAM5-MBP, TGA5-MBP, and WRKY53-MBP were transformed into *E. coli* Rosetta (DE3) cells for protein expression. MBP (as a control) was used in EMSAs with a LightShift Chemiluminescent EMSA Kit (Thermo Scientific) following the manufacturer’s protocol. The probe sequences used in the EMSA were as Supplementary 2.

### Firefly luciferase complementation imaging assay (LCI)

The *TGA5, CAM5*, and *CML8* coding sequences were cloned individually into the CLuc plasmid, and the *WRKY53* and *ACA8* coding sequences were cloned into the NLuc plasmid. The resulting plasmids were transformed into Agrobacterium strain GV3101. Luminescence signal from *Nicotiana tabacum* leaves infiltrated with the Agrobacterium cultures was imaged with a molecular imaging system (LB983, Berthold Technologies, Germany). Each leaf was divided into four quadrants prior to infiltration with the following combinations: *CLuc/NLuc, CAM5-CLuc/NLuc, CLuc/WRKY53-NLuc*, and *CAM5-CLuc/WRKY53-NLuc*; *CLuc/NLuc, CML8-CLuc/NLuc, CLuc/ACA8-NLuc*, and *CML8-CLuc/ACA8-NLuc*; or *CLuc/NLuc, TGA5-CLuc/NLuc, CLuc/WRKY53-NLuc*, and *TGA5-CLuc/WRKY53-NLuc*.

### *In vitro* pull-down assay

The coding sequences for *TGA5, CML8*, and *CAM5* were cloned into pGEX-4T-1, and the *ACA8* (aa 1–180), and *WRKY53* coding sequences were cloned into pET28A. The resulting CAM5-GST, CML8-GST, *TGA5*-GST, ACA8-HIS, and WRKY53-HIS plasmids were transformed into *E. coli* Rosetta (DE3) cells for protein production. Recombinant protein-GST was used as bait. The binding of CAM5-GST to WRKY53-HIS, *TGA5*-GST to WRKY53-HIS, and CML8-HIS to ACA8-GST was detected by immunoblot analysis using anti-GST and anti-HIS antibodies.

### Yeast two-hybrid assay

The coding sequences of *CML8* and *CAM5* were cloned into pGADT7, and the sequences encoding residues 1–217 of WRKY53 (WRKY53 [aa 1–217]) and residues 1–180 of ACA8 (ACA8 [aa 1–180]) were cloned into pGBKT7. The appropriate plasmid pairs CAM5-AD plus WRKY53 (1–217 aa)-BK and CML8-AD plus ACA8 (1–180 aa)-BK were co-transformed into competent yeast strain AH109 cells. The cells were grown on synthetic defined (SD) –Leu –Trp medium. After 4 days, growing colonies were transferred to SD medium –Ade –His –Leu –Trp for verification. Growing colonies on SD medium –Ade –His –Leu –Trp were transferred to SDmedium –Ade –His –Leu –Trp with X-α-gal. Cell growth on SD –Leu –Trp and SD –Ade –His – Leu –Trp and blue colonies on SD –Ade –His –Leu –Trp with X-α-gal indicate protein–protein interactions between the two proteins.

### Bimolecular fluorescence complementation assay

TGA5 was cloned into the pSPYNE vector via BamHI/SalI, WRKY53 was cloned into the pSPYCE vector via BamHI/SalI, and the plasmids were transformed into Agrobacterium strain GV3101. Both constructs (DORN1-YNE and RBOHF-YCE) were transformed into *N. tabacum* leaves infected with Agrobacterium. YFP fluorescence was imaged with confocal laser scanning microscopy (Leica TCS SP8, Leica Microsystems, Wetzlar, Germany) at an excitation wavelength of 510 nm and an emission wavelength of 510–530 nm.

### Statistical Analysis

Dunnett’s C (variance not neat) at the level of p < 0.05 was significant. Error bars denote ± standard error of mean (SEM).

## Results

### The evk-mediated ROS burst is dependent on RBOHD and RBOHF

We performed confocal laser scanning microscopy to measure H_2_O_2_ contents in Arabidopsis mesophyll cells. After 15 min of evk treatment, fluorescence from oxidized 2′,7′-dichlorofluorescein (DCF, reflecting H_2_O_2_ contents) consistently increased in mesophyll cells compared to the control (Figure 1a). Indeed, compared to the control, the fluorescence intensity increased by 196.2% (after 3 min), 309.1% (6 min), 252.9% (9 min), and 175.0% (15 min), suggesting that evk might cause mesophyll cells to generate H_2_O_2_ on a continual basis. The temporary addition of evk considerably reduced the fluorescence intensity of H_2_O_2_ in the NADPH oxidase *rbohd rbohf* double mutant. These findings suggest that the evk-induced ROS burst is mediated by NADPH oxidase. The fluorescence intensity after evk treatment rose by 268.9% (3 min), 285.5% (6 min), 217.4% (9 min), and 199.8% (15 min) in wild-type plants pretreated with ruthenium red (a Ca^2+^ channel inhibitor that blocks the release of internal Ca^2+^), values that were not significantly different from those of the WT+evk treatment group (Figure 1b). These results demonstrate that NADPHase is involved in the evk-induced ROS burst and that changes in calcium concentrations have no effect on the evk-induced H_2_O_2_ burst.

**Figure 1.**
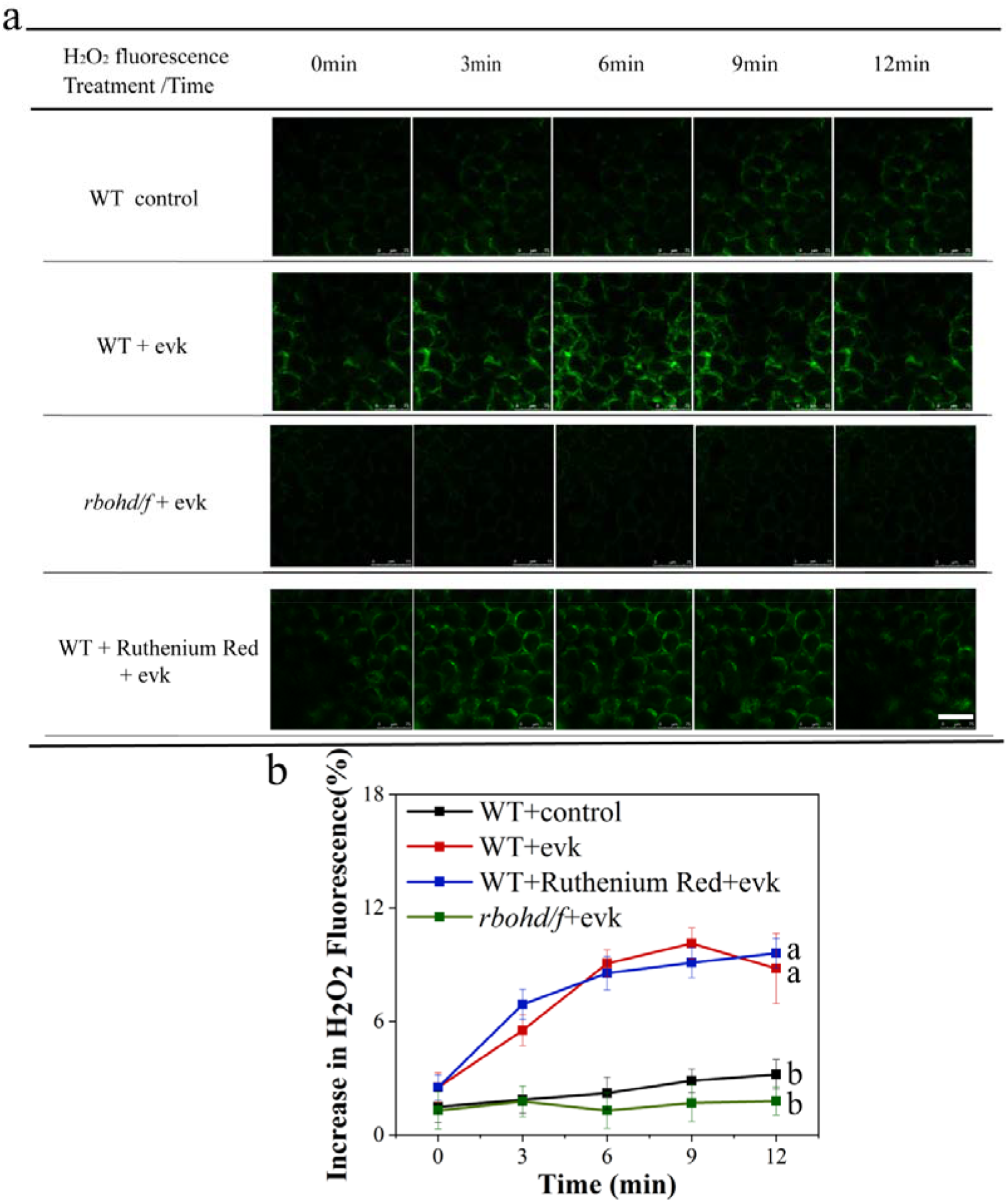
evk-induced increases in H_2_O_2_ levels depend on RBOHD, RBOHF but not on internal Ca^2+^ release channels in Arabidopsis mesophyll cells. (a) ROS burst in response to evk treatment in the wild type (WT), *rbohd rbohf*, and WT pretreated with ruthenium red. The cells were washed to remove the 50 μM H_2_DCF-DA test solution (0 min), evk was added, and the cells were photographed every 3 min. Scale bars, 75 μm. (b) H_2_O_2_ fluorescence over time in ruthenium red–treated and *rbohd rbohf* cells compared to the WT evk group. Analysis of changes in H_2_O_2_ fluorescence indicated that increases in H_2_O_2_ fluorescence were significantly suppressed in *rbohd rbohf* cells compared to the WT + evk-treated group but were unaffected in ruthenium red–treated cells. The seedlings plants were grown for 10 days. Error bars denote ± SEM, n ≥ 18, and means labeled with different letters are significantly different at p < 0.05, Dunnett’s C (variance not neat).

### Docking simulations of evk and RBOHD/RBOHF

Our experimental results indicated that the evk-mediated ROS burst is dependent on RBOHD and RBOHF, but how evk interacts with RBOHD/RBOHF is still unknown. Therefore, docking simulation was carried out to explore their binding mode. The 3D structures of RBOHD/RBOHF were built with SwissModel firstly. And then the possible binding sites of RBOHD/RBOHF were explored with Site Finder in Molecular Operating Environment (MOE). The top two recommended binding sites were used as binding sites of RBOHD/RBOHF, which are corresponding to the FAD (site1) and/or NADPH binding site (site2) of the highly homologous human DUOX1-DUOXA1 (dual oxidase 1 and dual oxidase maturation factor 1)^63^ and Cylindrospermum stagnale NADPH oxidase 5 (NOX5)^64^. In the NADPH oxidase, NADPH works as an electron donor and FAD as an electron transporter, which passes electron sequentially to the two heme molecules and then to oxygen molecule, and forms superoxide anion finally^65^. So evk was predicted inducing ROS production by assisting electron transfer, their possible binding modes were shown in figure 2.

**Figure 2.**
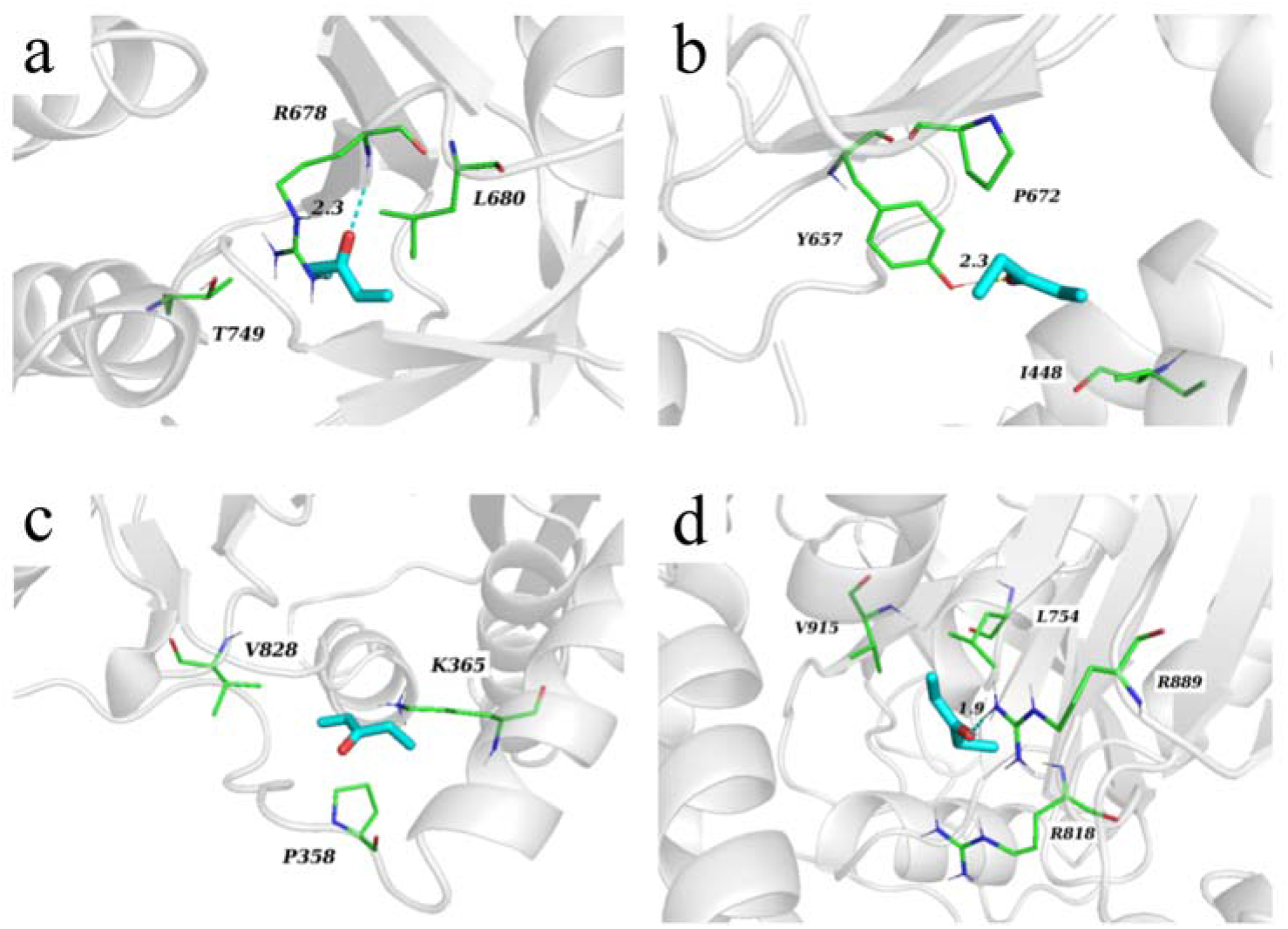
The structural docking model of evk (cyan sticks) binding on site1 (a) or site2 (c) of RBOHD, site1 (b) or site2 (d) of RBOHF (white cartoon) and, and the key residues labelled out in green sticks, and the hydrogen bond interactions labelled out in dash line.

As shown in Fig. 2a, evk bound to site1 of RBOHD by forming hydrogen bond interaction with Arg-678, hydrophorbic interactions with Leu-680, Thr-749. And evk bound to site1 of RBOHF (Fig. 2b) by forming hydrogen bond interaction with Tyr-657, hydrophorbic interactions with Pro-672, Ile-448. Besides, evk can bind with RBOHD on site2 (Fig. 2c) by forming hydrophorbic interaction with Pro-358, Lys-365, Val-828; or bound with RBOHF on site2 (Fig. 2d) by forming hydrogen bond interactions with Arg-889, forming hydrophorbic interactions with Leu-754, Val-915. Based on the docking simulation results, evk was predicted to assist electrons transfer of RBOHD/RBOHF by binding to their FAD or NADPH binding site, and thus inducing ROS production.

### RBOH participates in the evk-induced increase in intracellular calcium concentrations

We used confocal laser scanning microscopy to detect intracellular Ca^2+^ levels based on Fluo3-AM fluorescence intensity. Compared to the control, the fluorescence intensity of Fluo3-AM increased by 183.9% (20 s), 189.2% (40 s), 132.1% (60 s), 124.3% (80 s), and 119.4% (100 s) into evk treatment. Ca^2+^ fluorescence was substantially lower in the WT pretreated with ruthenium red and *rbohd rbohf* cells plus evk than in WT cells exposed to evk (Figure 3a). We determined that the intracellular calcium pool is the source of the evk-induced calcium signal, as the fluorescence intensity of intracellular Ca^2+^ Fluo3-AM decreased by 118.6% in *rbohd rbohf* compared to the control after 40 s (Figure 3b). These results indicate that NADPHase affects the evk-induced calcium burst, whereas the evk-mediated H_2_O_2_ burst is unaffected by intracellular calcium pool inhibitors, indicating that H_2_O_2_ functions upstream of calcium signaling.

**Figure 3.**
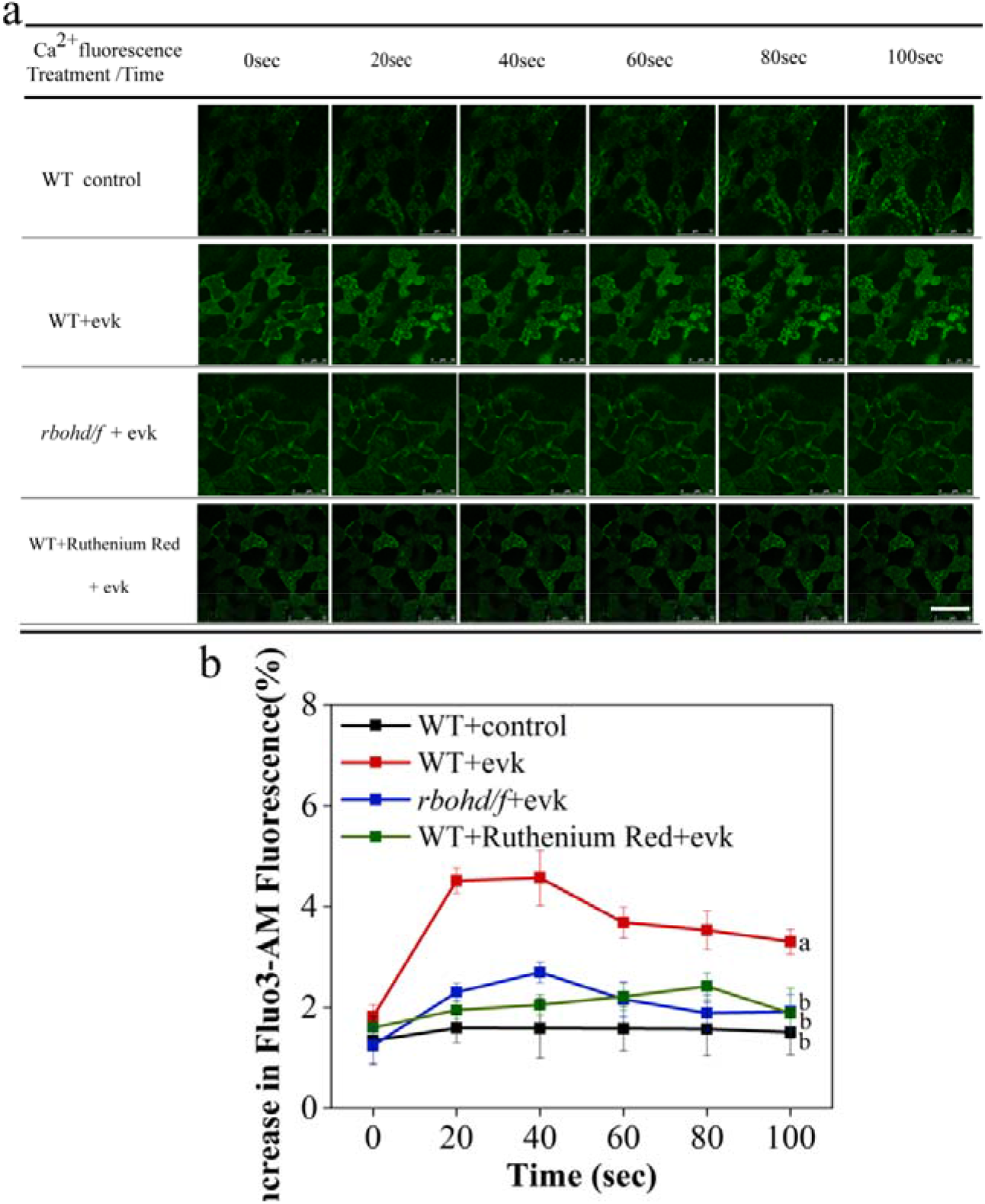
evk-induced increases in intracellular calcium levels depend on RBOHD RBOHF and internal Ca^2+^ release channels in Arabidopsis mesophyll cells. (a) The cells were washed to remove the 10 μM Fluo3-AM test solution (0 min), evk was added, and the cells were photographed every 30 s. Scale bars, 50 μm. (b) evk-induced increases in Ca^2+^ fluorescence are blocked in ruthenium red–treated WT and *rbohd rbohf* cells compared to WT + evk cells. The seedlings were grown for 10 days. Error bars denote ± SEM, n ≥ 19, and means labeled with different letters are significantly different at p < 0.05, Dunnett’s C (variance not neat).

### RBOH participates in evk-mediated calcium efflux

NMT revealed that treating Arabidopsis mesophyll cells with transient evk results in an intense efflux of transmembrane calcium ions from mesophyll cells. The efflux flow rate reached 261.11 ± 20.57 pmol cm^−2^ s^−1^ and then gradually returned to the resting state. To explore the role of NADPHase in Ca^2+^ efflux induced by evk, we examined the NADPHase mutant *rbohd rbohf* to detect changes in Ca^2+^ flow after evk treatment. Evk-induced Ca^2+^ efflux was weaker in *rbohd rbohf* compared to WT (Figure 4a). Immediately after the addition of evk, the calcium ion efflux flow rate was 111.83 ± 24.11 pmol cm^−2^ s^−1^. The peak calcium efflux in WT and *rbohd rbohf* plants was 237.19 ± 15.31 pmol cm^−2^ s^−1^ and 58.95 ± 16.37 pmol cm^−2^ s^−1^, respectively (Figure 4b), indicating that calcium efflux was significantly reduced in the mutant. These results indicate that RBOHD and RBOHF are involved in evk-induced calcium efflux in mesophyll cells.

**Figure 4.**
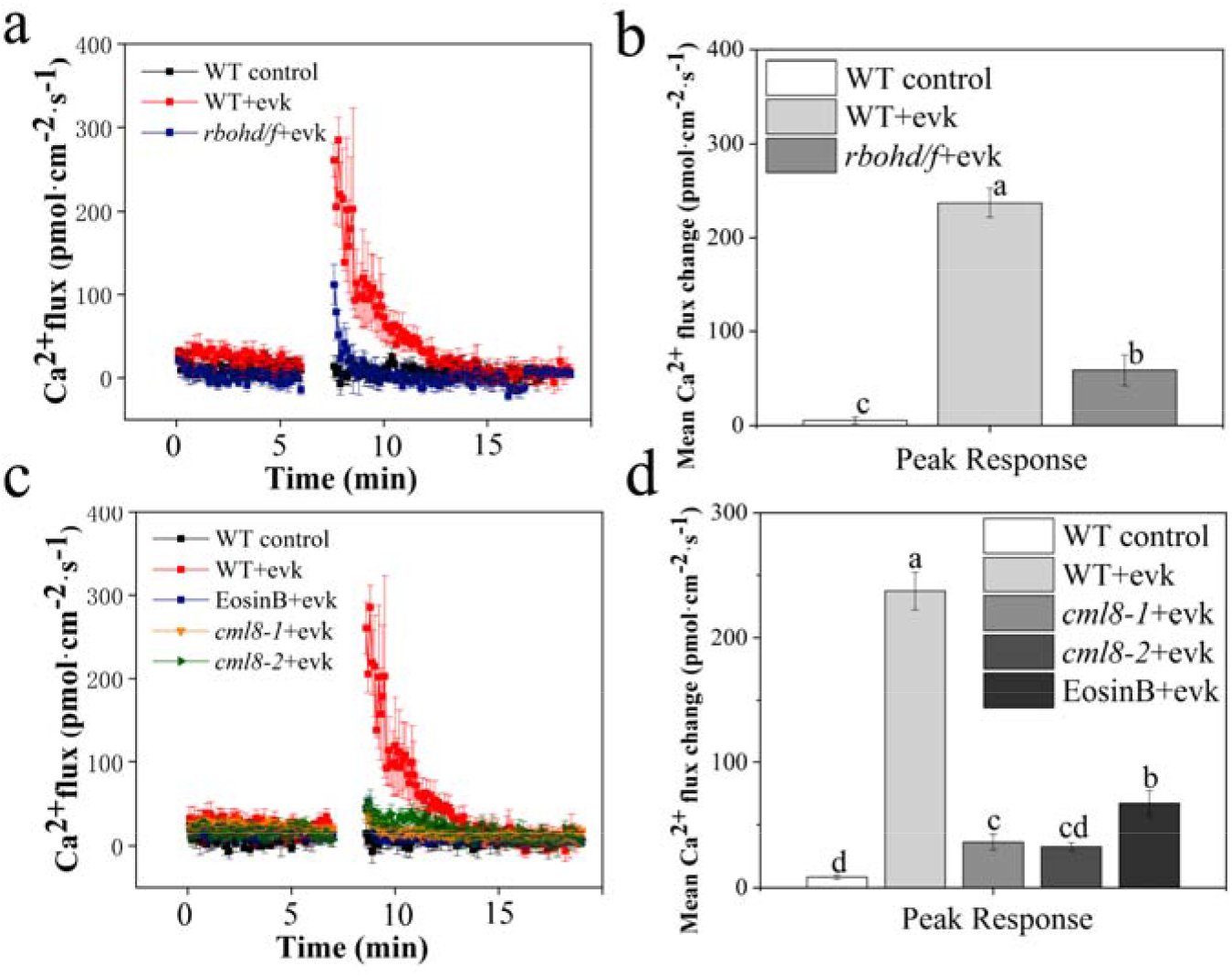
evk-induced Ca^2+^ efflux is enhanced by H_2_O_2_ and suppressed in eosin B–pretreated WT and *cml8* plants. (a,c) evk treatment increases Ca^2+^efflux in WT, but not in *rbohd rbohf, cml8-1*, or *cml8-2*. (b,d) Peaks in mean Ca^2+^ efflux among genotypes and treatments, showing that Ca^2+^ efflux is significantly suppressed in all groups compared to the evk-treated WT control. The plants were grown for 3 weeks. Error bars denote ± SEM, n ≥ 6, and columns labeled with different letters are significantly different at p < 0.05, Dunnett’s C (variance not neat).

### CML8 interacts with ACA8 to regulate evk-induced calcium efflux

Eosin B is an inhibitor of calcium pumps. We therefore pretreated mesophyll cells with 10 µM eosin B, followed by evk treatment. Calcium ion efflux was significantly lower upon treatment with eosin B, reaching a peak value of 102.39 ± 23.73 pmol cm^−2^ s^−1^ and a peak calcium efflux value of 67.28 ± 10.34 pmol cm^−2^ s^−1^, which was significantly lower than the peak value of the WT + evk group (237.19 ± 15.31 pmol cm^−2^ s^−1^), indicating that the calcium pump plays a role in evk-mediated calcium ion efflux from mesophyll cells. The *cml8-1* and *cml8-2* mutants also showed lower calcium ion efflux in mesophyll cells under evk treatment, with peak values of 44.27 ± 2.47 pmol cm^−2^ s^−1^ and 43.64 ± 12.48 pmol cm^−2^ s^−1^, respectively (Figure 4c). The peak calcium efflux values in *cml8-1* and *cml8-2* were 36.48 ± 6.21 pmol cm^−2^ s^−1^ and 32.85 ± 3.20 pmol cm^−2^ s^−1^, respectively, which were significantly lower than the peak value of the WT + evk group (Figure 4d). These results indicate that CML8 regulates calcium pump function during evk-mediated calcium ion efflux, thus affecting calcium efflux. We established that the calcium-dependent ATPase ACA8 is the likely target of CML8 regulation, as demonstrated by its interaction in yeast two-hybrid, firefly luciferase complementation imaging, and pull-down assays (Figure 5a-5c).

**Figure 5.**
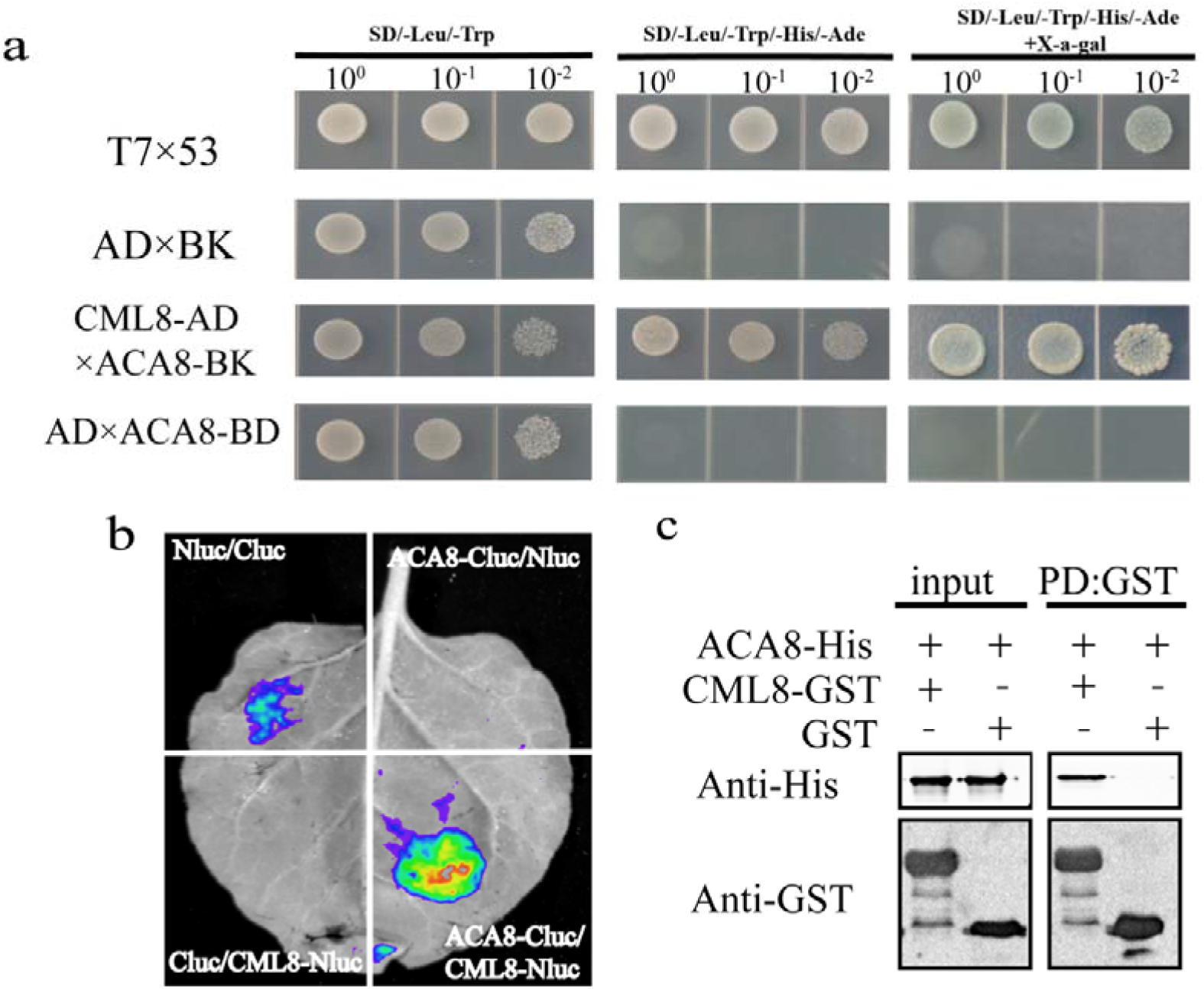
CML8 interacts with ACA8 (a) Yeast two-hybrid assay showing that ACA8 interacts with CML8. Yeast colonies harboring the indicated constructs were grown on synthetic defined (SD) medium lacking Trp and Leu, or SD medium lacking Trp, Leu, His, and Ade. (b) Firefly luciferase complementation imaging assay. *N. benthamiana* leaves were infiltrated with the construct pairs *ACA8-CLuc/CML8-NLuc, Cluc/NLuc, ACA8-CLuc/NLuc*, and *CLuc/CML8-NLuc*. (d) *In vitro* pull-down assays. The N terminus (residues 1–180), an intracellular domain, was used to interact with CML8. GST-CML8 pulled down His-tagged ACA8, indicating that CML8 interacts with ACA8.

### Evk mediates SA/JA antagonism in Arabidopsis

Evk activates the H_2_O_2_ burst. H_2_O_2_ and SA are strongly correlated, and *WRKY53* expression is strongly activated by evk, as revealed by reverse transcription quantitative PCR (RT-qPCR). Therefore, we reasoned that evk might activate SA-related gene expression. We measured the expression of SA-related genes via RT-qPCR. We observed that evk treatment upregulates *NPR1, NPR4, ICS1* (*ISOCHORISMATE SYNTHASE 1*), *PAD4* (*PHYTOALEXIN DEFICIENT 4*), and *SARD1* (*SAR DEFICIENT 1*), especially during the early stage of treatment (0 to 3 h). However, *PR1* was not only upregulated at 3 h, but it was still upregulated at 8 h, indicating that a regulatory substance activated *PR1* expression during the early and late stages of treatment. Most SA-related genes were downregulated in the later stage of treatment, suggesting that JA might antagonize their expression at this stage. However, the JA*-*related genes *OPR1* (*12-OXOPHYTODIENOATE REDUCTASE 1*), *OPR3, AOC3* (*ALLENE OXIDE CYCLASE 3*), *JMT* (*JASMONIC ACID CARBOXYL METHYLTRANSFERASE*), *LOX2*, and *PDF1*.*2* were strongly upregulated during the later stage of evk treatment (3 to 8 h). RT-qPCR showed that evk treatment antagonizes SA/JA signaling, especially the simultaneous activation of *PR1* and *PDF1*.*2* expression at 8 h. The finding that *PR1* and *PDF1*.*2* are both upregulated at the later stage of treatment (Figure 6a,6b) is intriguing. In addition, the plants accumulated more SA within 3 h of evk treatment. During the later stage of treatment (8 h), the SA content remained high, and the JA level increased (Figure 6c).

**Figure 6.**
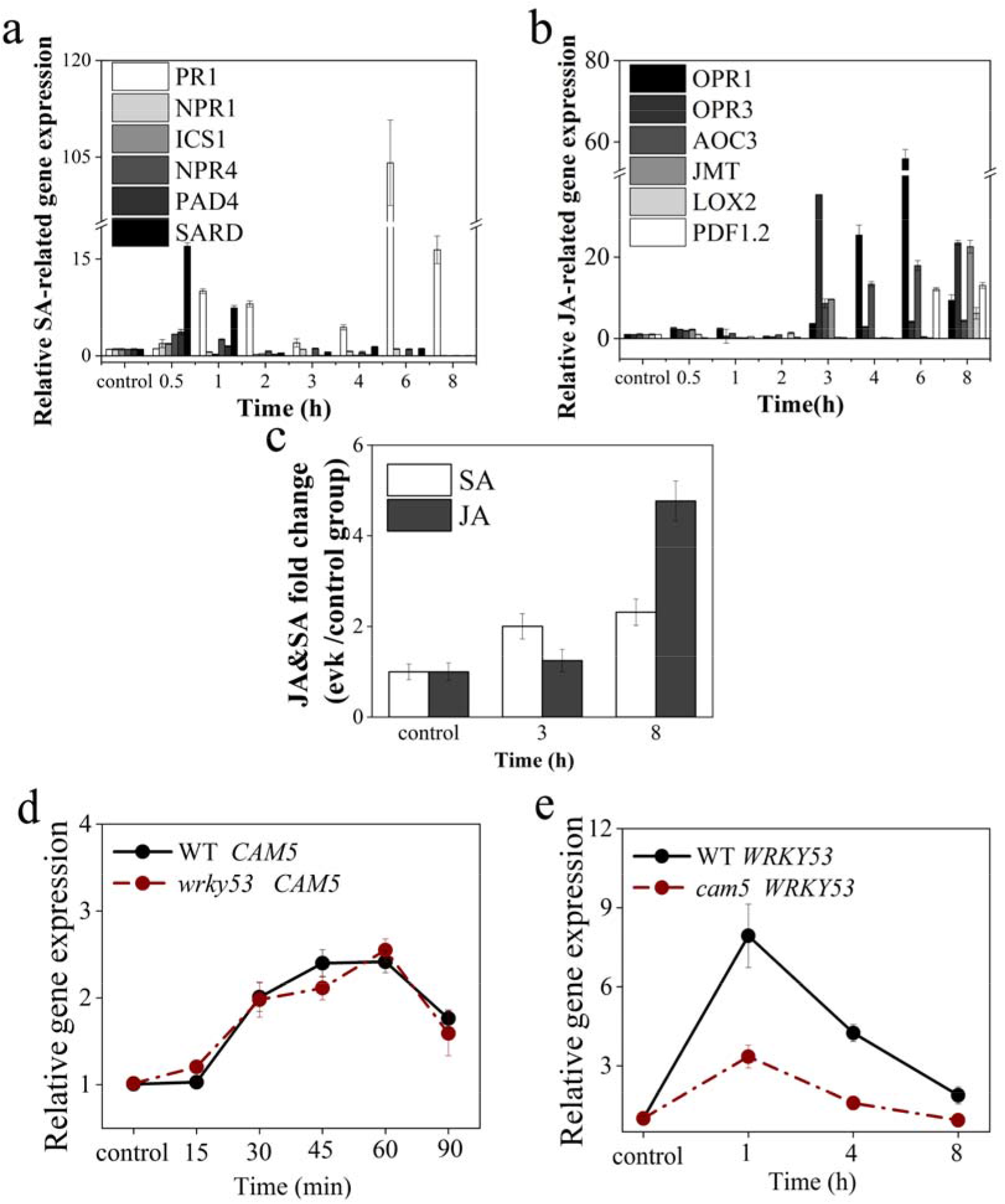
Evk-induced changes in gene expression. (a,b) Expression levels of key SA/JA-related genes in WT plants following evk treatment. The plants were grown for 3 weeks. Data are means ± SEM, n ≥ 3. (c) evk-induced fold-change in JA and SA levels. (d) Relative *CAM5* expression levels in the leaves of 2-week-old WT and *wrky53* seedlings. (e) Relative *WRKY53* expression levels in the leaves of 2-week-old WT and *cam5* seedlings. The seedlings were grown for 3 weeks. Data are means ± SEM, n ≥ 3.

### CAM5 functions upstream of WRKY53 in evk signaling

RT-qPCR analysis showed that *WRKY53* expression begins to decline after 1 h of upregulation (Figure 6e). We examined *CAM5* expression during early evk treatment (90 min) because the calcium signal is an early signal. *CAM5* was upregulated at all time points during the early stage of evk treatment (Figure 6d). We also measured *WRKY53* expression in the *cam5* mutant. *WRKY53* expression was much lower in the mutant. On the contrary, *CAM5* expression was not affected in the *wrky53* mutant (Figure 6d,6e). These results indicate that CAM5 functions upstream of WRKY53 in response to evk treatment.

### CAM5 and WRKY53 positively regulate the expression of SA-related genes and negatively regulate the expression of JA-related genes

Within 3 h of evk treatment, *PR1, NPR4*, and *SARD* were significantly upregulated in the WT but significantly downregulated in *wrky53* and *cam5* (Figure 7e,7f,7g), indicating that WRKY53 and CAM5 positively regulate the evk-induced activation of *PR1, NPR4*, and *SARD1*. After 3 h of evk treatment, JA-related genes were significantly upregulated, especially *OPR1, OPR3, JMT*, and *AOC3* (Figure 7a,7b,7c,7d). Notably, in *wrky53*, the upregulation of these genes was more obvious, indicating that WRKY53 negatively regulates JA-related gene expression. When *WRKY53* was mutated, we detected little evk-induced upregulation of *PR1*. On the contrary, *PDF1*.*2* was significantly upregulated during the late stage of evk treatment, indicating that WRKY53 is likely a positive regulator of *PR1* and a negative regulator of *PDF1*.*2* (Figure 7h,7i).

**Figure 7.**
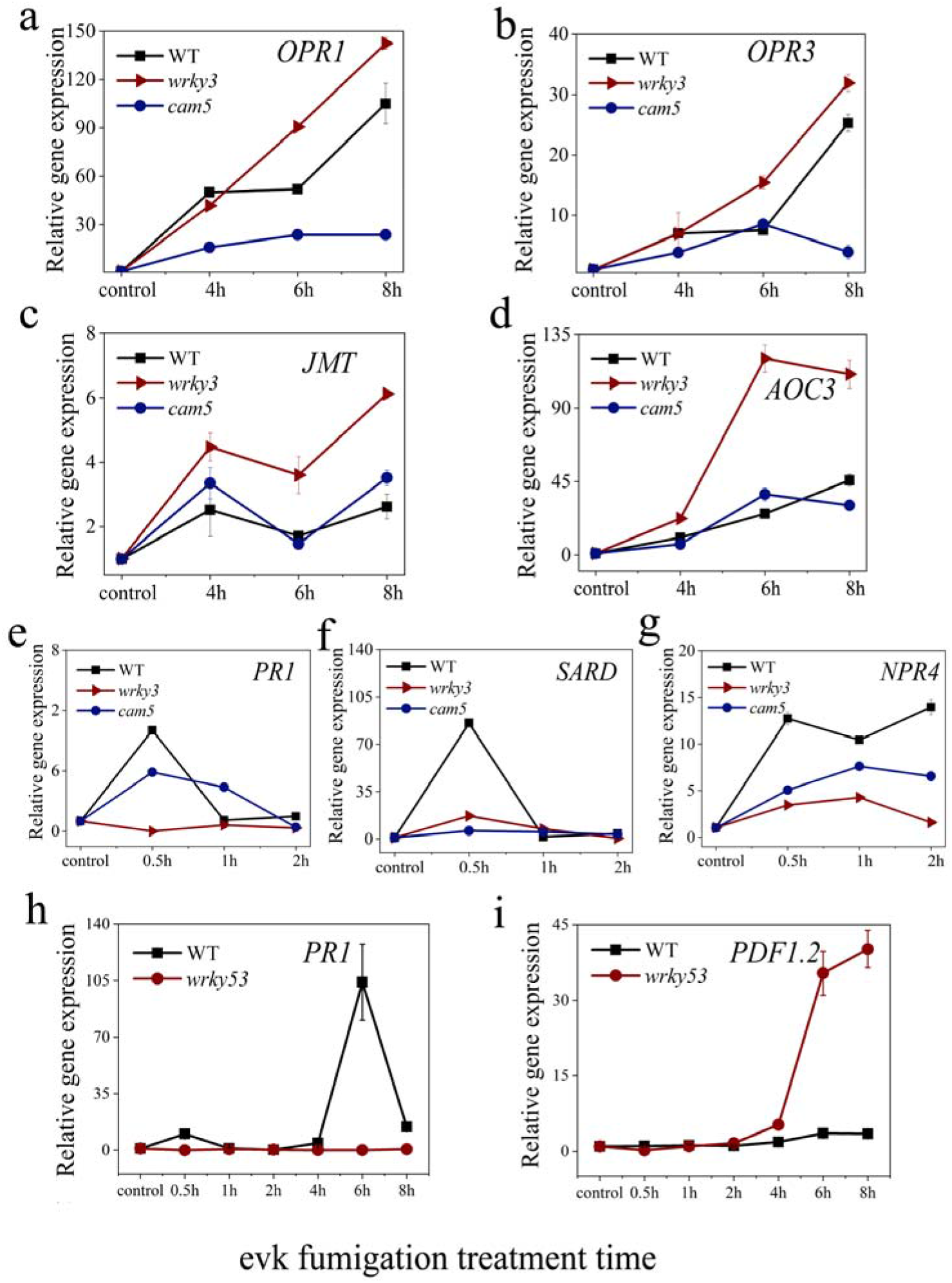
Evk-induced changes in SA/JA-related gene expression in WT, *wrky53*, and *cam5* plants (a-d) Relative expression levels of JA-related genes in the leaves of WT, *wrky53*, and *cam5* plants following evk treatment. (e-g) Relative expression levels of SA-related genes in the leaves of WT, *wrky53*, and *cam5* plants following evk treatment. (h,i) Relative expression levels of *PR1* and *PDF1*.*2* in the leaves of WT and *wrky53* plants following evk treatment. The plants were grown for 3 weeks. Data are means ± SEM, n ≥ 3.

### CAM5 interacts with WRKY53, and increasing calcium concentrations weaken this interaction

Evk treatment upregulated the expression of SA-related genes, which was strongly associated with both WRKY53 and CAM5, and CAM5 regulates the expression of *WRKY53*, suggesting that WRKY53 and CAM5 might interact. In yeast two-hybrid assays, yeast cells harboring CAM5-AD and WRKY53-BD (amino acids [aa] 1–217) grew well on selective medium, and the colonies turned blue on X-gal plates, indicating that these two proteins indeed physically interact (Figure 8a). In a luciferase complementation test (Figure 8c), *Nicotiana benthamiana* leaves transiently expressing *CAM5-NLuc* + *WRKY53-CLuc* constructs had stronger luminescence than the *NLuc + CLuc, CAM5-NLuc + CLuc*, or *WRKY53-CLuc + NLuc* combinations (Figure 8c), confirming that CAM5 and WRKY53 interact. Finally, in a glutathione S-transferase (GST) pull-down experiment, recombinant CAM5-GST pulled down WRKY53-His (Figure 7d). In response to increasing calcium ion concentrations, the amount of WRKY53 pulled down by CAM5-GST gradually decreased. In the presence of EGTA, CAM5-GST pulled down the greatest amount of WRKY53 (Figure 8b). These results indicate that WRKY53 is released from the CAM5-WRKY53 complex in the presence of increasing calcium concentrations.

**Figure 8.**
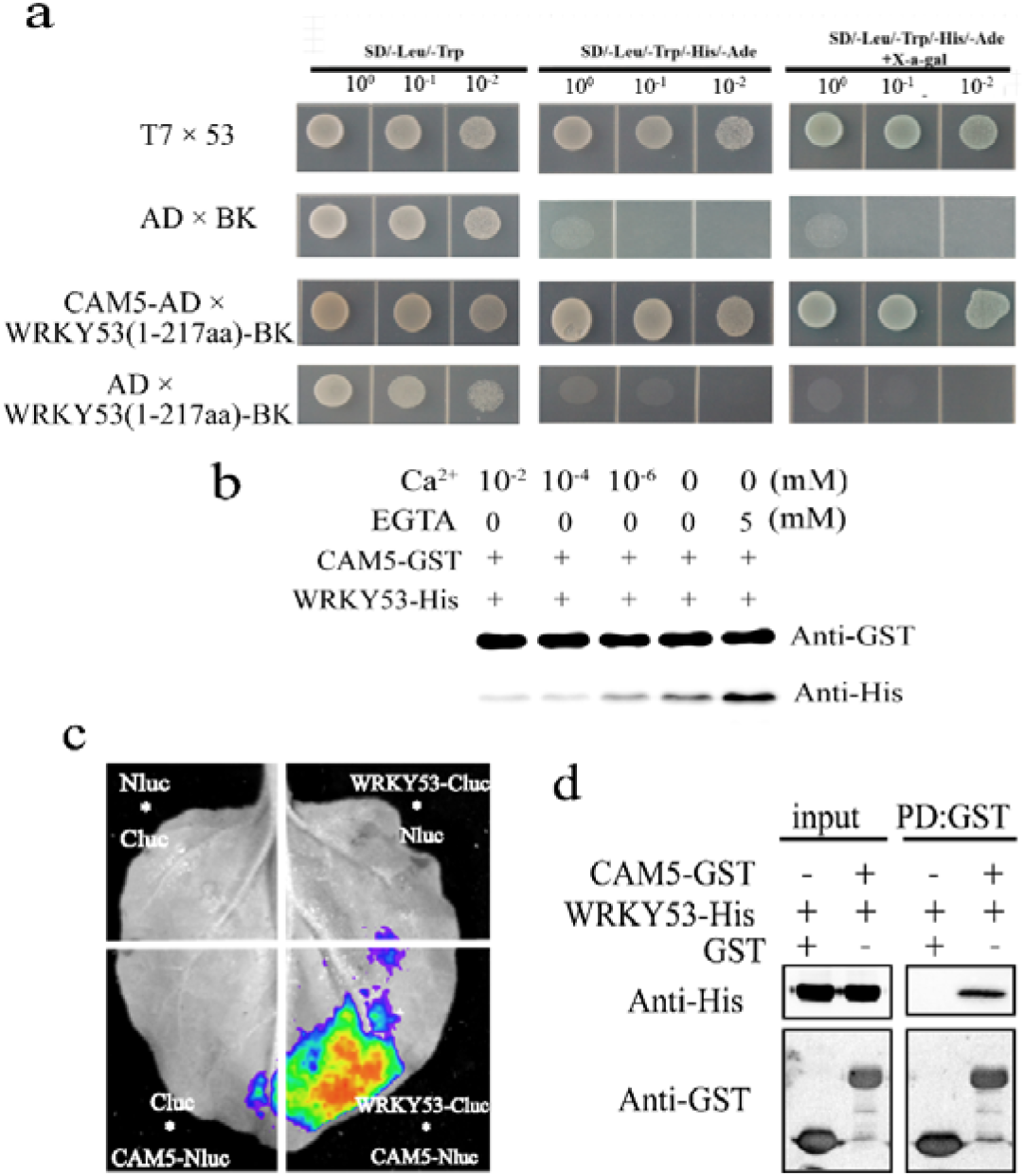
CAM5 interacts with WRKY53 (a) Yeast two-hybrid assay; the N terminus of WRKY53 (residues 1–217) was used as bait. Yeast cells harboring CAM5-AD and WRKY53-BD (aa 1–217) grew normally on all plates. (b) Pull-down assay showing the effects of calcium concentration on the binding of WRKY53 to CAM5. (c) LCI assay showing the interaction between WRKY53 and CAM5. *N. benthamiana* leaves were infiltrated with the pairs of constructs *WRKY53-CLuc/CAM5-NLuc, CLuc/NLuc, WRKY53-CLuc/NLuc*, or *CLuc/CAM5-NLuc*. (d) GST pull-down assay showing that GST-CAM5 binds to WRKY53-His.

### TGA5 interacts with WRKY53

We hypothesized that TGA5, a transcription factor that responds to RES, might be involved in the late stage of the response to evk treatment by upregulating *PR1* after the downregulation of *WRKY53*. In a luciferase complementation test, we detected higher luciferase activity from *N. benthamiana* leaves co-infiltrated with the constructs *TGA5-CLuc + WRKY53-NLuc* than with the pairs *NLuc + CLuc, TGA5-CLuc + NLuc*, and *WRKY53-NLuc* + *CLuc* (Figure 9b), indicating that TGA5 and WRKY53 interact. In a GST pull-down assay, recombinant TGA5-GST pulled down WRKY53-His (Figure 9a). We also established that TGA5 interacts with WRKY53 in the nucleus, as demonstrated by BiFC (Figure 9c). These results demonstrate that TGA5 and WRKY53 interact *in vitro* and *in vivo*.

**Figure 9.**
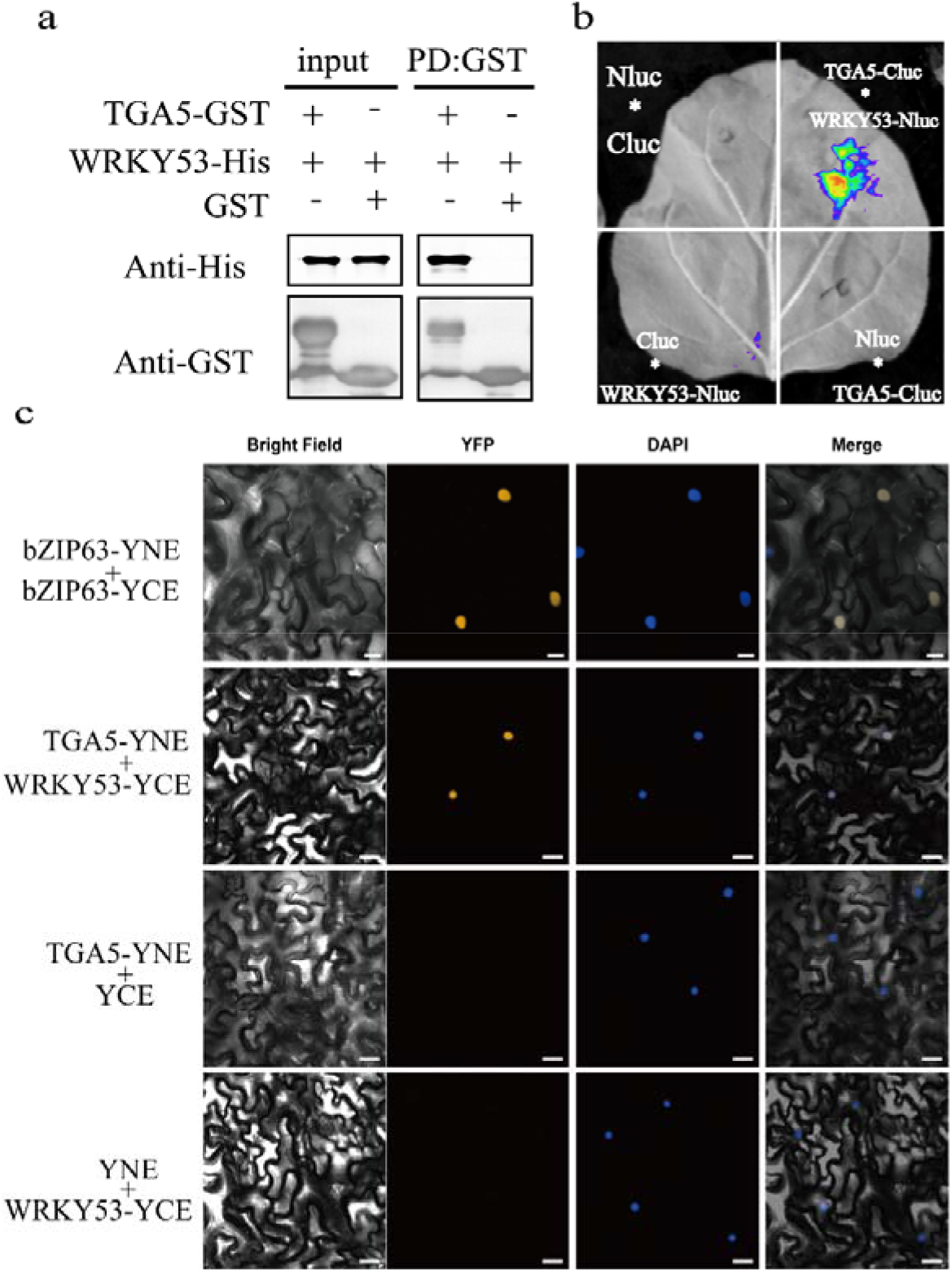
WRKY53 interacts with TGA5 (a) TGA5-GST pulls down WRKY53-His *in vitro*. (b) LCI assay showing that WRKY53 and CAM5 interact *in vivo. N. benthamiana* leaves were infiltrated with the pairs of constructs *WRKY53-CLuc/CAM5-NLuc, CLuc/NLuc, WRKY53-CLuc/NLuc*, and *CLuc/CAM5-NLuc*. (c) BiFC assays indicating that TGA5 interacts with WRKY53 in the nucleus.

### WRKY53 enhances the binding of TGA5 to *PR1* and weakens its binding to *PDF1*.*2*

We performed EMSAs to test whether WRKY53 might bind to the *PR1* promoter. Accordingly, we labeled a promoter fragment of *PR1* with biotin and incubated the resulting probe with recombinant purified His-WRKY53. We observed that WRKY53 directly binds to the w-box in the *PR1* promoter, whereas CAM5 did not (Figure 10a). In the presence of both WRKY53 and CAM5, the binding of WRKY53 to the w-box of the *PR1* promoter was weakened. When calcium ions were added to the system, the binding of WRKY53 to the w-box was restored (Figure 10a). To determine whether TGA5 also bound to the w-box in the *PR1* promoter, we performed another EMSA with recombinant TGA5-MBP (maltose-binding protein), finding that TGA5 directly binds to the w-box in the *PR1* promoter. In the presence of both WRKY53 and TGA5, WRKY53 appeared to enhance the binding strength of TGA5 to this w-box. These results indicate that WRKY53 promotes the binding of TGA5 to the w-box in the *PR1* promoter (Figure 10b). Moreover, WRKY53 bound to the w-box in the *PDF1*.*2* promoter, whereas CAM5 did not (Figure 10c). In the presence of both WRKY53 and CAM5, the binding of WRKY53 to the w-box in the *PDF1*.*2* promoter was weakened. When calcium ions were added to this system, the original binding strength of WRKY53 to the w-box in the *PDF1*.*2* promoter was restored (Figure 10c). WRKY53 failed to bind to the as-1 sequence in the *PDF1*.*2* promoter, but TGA5 bound to this sequence. In the presence of both WRKY53 and TGA5, WRKY53 weakened the binding of TGA5 to the as-1 sequence (Figure 10d). In summary, when the calcium ion concentration increases, WRKY53 is released from the CAM5-WRKY53 complex, allowing it to interact with the downstream protein TGA5 to jointly regulate the expression of *PR1* and *PDF1*.*2*. WRKY53 enhances the binding strength of TGA5 to the w-box in the *PR1* promoter, while WRKY53 weakens the binding strength of TGA5 to as-1 in the *PDF1*.*2* promoter.

**Figure 10.**
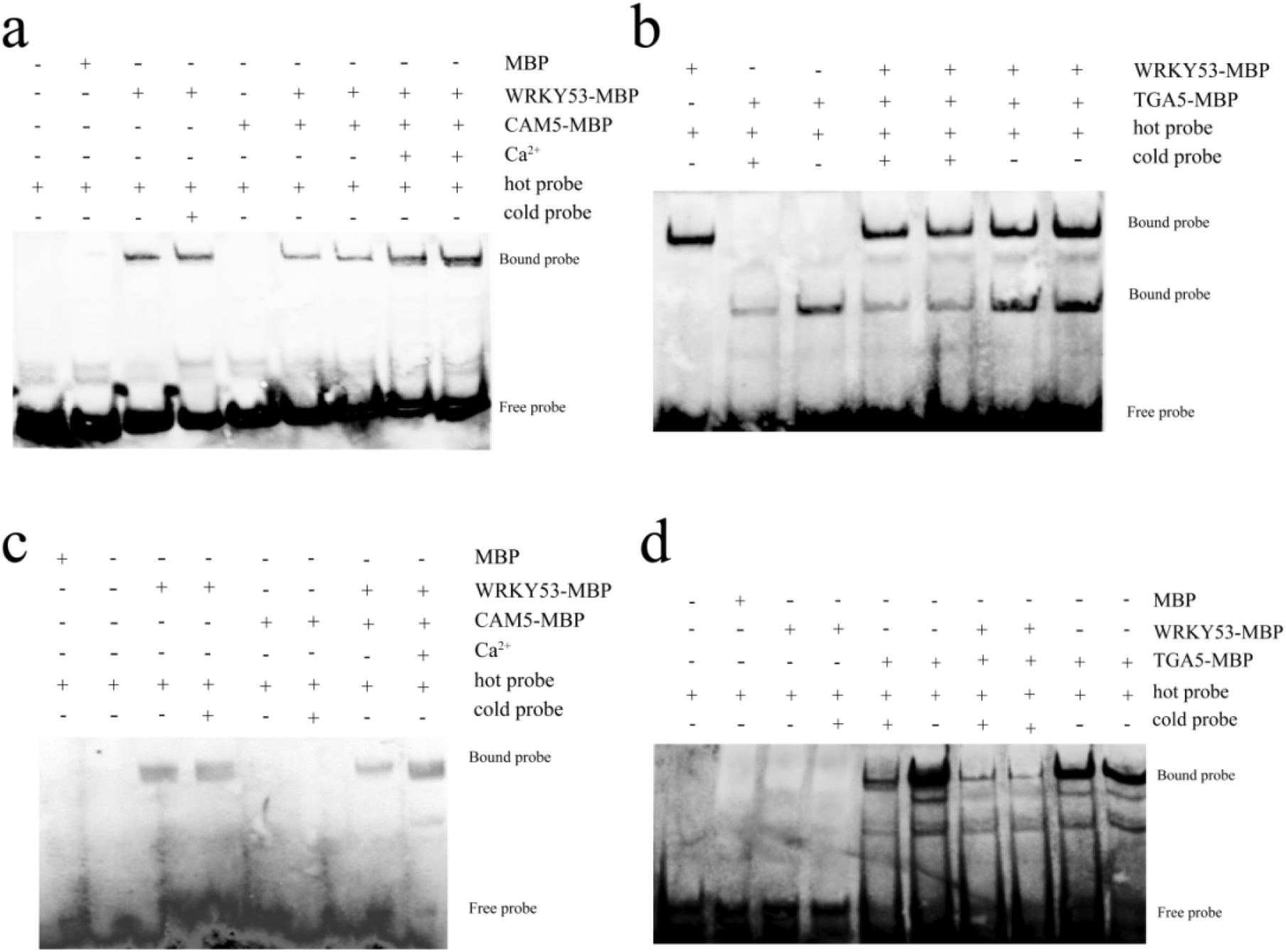
WRKY53 enhances the binding strength of TGA5 to the w-box in the *PR1* promoter, while WRKY53 weakens the binding strength of TGA5 to as-1 in the *PDF1*.*2* promoter (a) Ca^2+^ and CAM5 influence the binding ability of recombinant WRKY53 to the *PR1* promoter, as determined by EMSA. Hot probe refers to biotin-labeled probe and cold probe to unlabeled probe. The Ca^2+^ concentration is 10 mM. (b) WRKY53 enhances the binding of TGA5 to the w-box in the *PR1* promoter. WRKY53 binds to the w-box sequence ((C/T)TGAC(T/C)) in the *PR1* promoter. TGA5 binds to the w-box in the *PR1* promoter. WRKY53 enhances the binding of TGA5 to the w-box in the *PR1* promoter. The w-box sequence (C/T)TGAC(T/C) overlaps with the action center element as-1 (TGAC) of TGA. (c) Ca^2+^ and CAM5 influence the binding of WRKY53 to the *PDF1*.*2* promoter. (d) WRKY53 weakens the binding of TGA5 to the as-1 sequence in the *PDF1*.*2* promoter.

### Dual-luciferase reporter assay

Finally, to verify that WRKY53 and TGA5 activate the promoters of their downstream genes, we performed transient expression assays in Arabidopsis protoplasts and *N. benthamiana* leaves. To this end, we cloned the *PR1* promoter into pGreenII0800-LUC to generate the luciferase (LUC) reporter construct. In parallel, we cloned the coding sequences of *WRKY53* and *TGA5* in the pGreenII 62-SK vector to generate the effector constructs, using *35S:REN* as an internal control for transfection or infiltration efficiency. The luciferase activation assay showed that WRKY53 and TGA5 induce *PR1* transcription, while only TGA5 induced *PDF1*.*2* transcription, and only WRKY53 repressed *PDF1*.*2* transcription. We then examined the activity of the effectors *WRKY53* and *TGA5* in regulating the *PR1* and *PDF1*.*2* promoters when co-expressed. We observed that WRKY53 cooperates with TGA5 to positively regulate *PR1* transcription (Figure 11c, 11d). Similarly, we determined that TGA5 inhibits the induction of *PDF1*.*2* transcription mediated by WRKY53 (Figure 11e, 11f).

**Figure 11.**
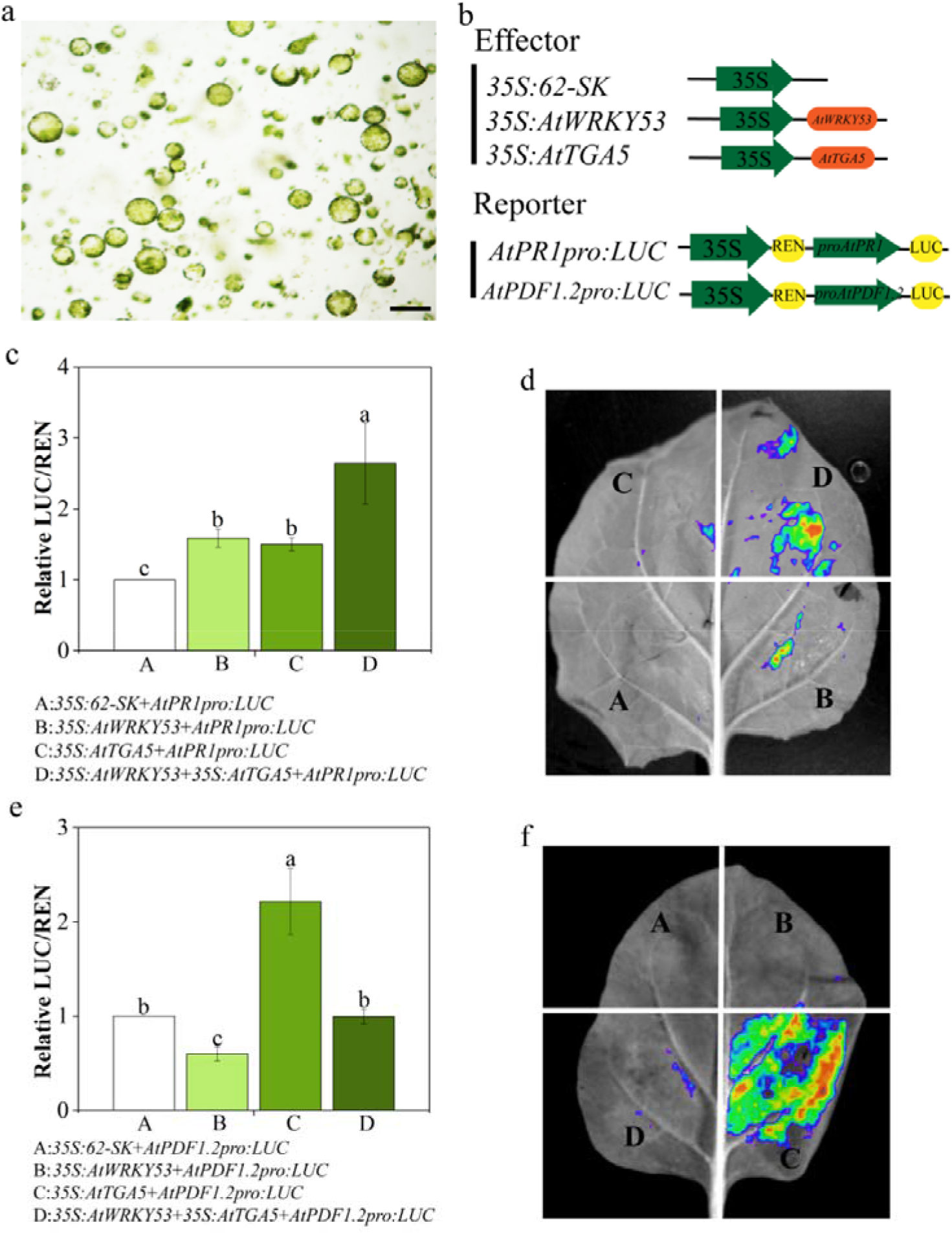
Luciferase activation assay. (a) Protoplasts extracted from 5-week-old Arabidopsis leaves. (b) Schematic diagrams of the effector and reporter constructs used for the dual-luciferase assay. (c,e) Dual-luciferase assay. Arabidopsis protoplasts were transiently transfected with the indicated constructs. Relative luciferase activity is shown as the ratio of LUC to REN activity. *35S:REN* (in pGreenII 62-SK) was used as an internal control. Error bars denote ± SEM, n ≥ 16, and means labeled with different letters are significantly different at p < 0.05, Dunnett’s C (variance not neat). (d,f) Transient infiltration assay of *N. benthamiana* leaves with the indicated constructs.

**Figure 12.**
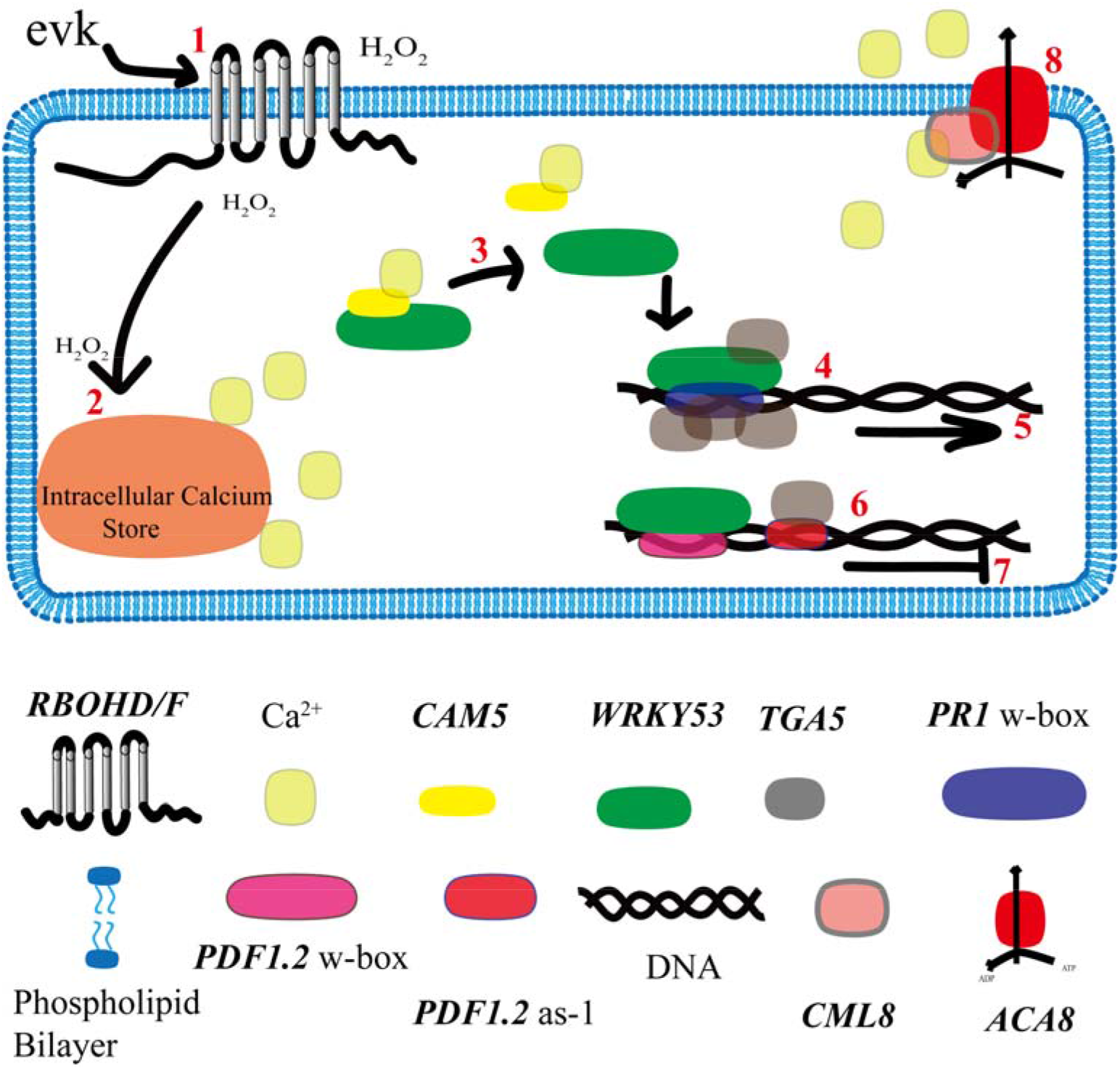
Schematic diagram of plant responses to evk.(1) evk is recognized by plants and causes ROS release. (2) H_2_O_2_ mediates the release of calcium ions from the intracellular calcium pool and increases the intracellular calcium concentration. (3) The WRKY53-CAM5 complex is disrupted under high calcium concentrations. (4) WRKY53 binds to the w-box in the *PR1* promoter to induce its expression, and (5) WRKY53 enhances the binding of TGA5 to as-1 in the *PR1* promoter to enhance its expression. (6) WRKY53 binds to the w-box in the *PDF1*.*2* promoter to inhibit its expression, and (7) WRKY53 weakens the binding of TGA5 to as-1 in the *PDF7* promoter, thereby inhibiting its expression. When *WRKY53* expression decreases during the later period of evk treatment (8 h), the inhibition of *PDF1*.*2* is alleviated, thus significantly increasing *PDF1*.*2* expression. (8) Calcium ions are expelled from the cell by ACA8-CML8, and calcium ion levels return to the resting state.

## Discussion

When a herbivore damages the leaves of a plant, volatile compounds such as terpenoids, C6-volatiles, and electrophiles are released. These compounds elicit a range of defense mechanisms in both the damaged plant and nearby plants. Infestation with lima bean mite (*Tetranychus urticae*) causes the release of terpenoid compounds, which in turn elicits the expression of defense genes in nearby plants (Arimura et al., 2000)^66^. These volatiles serve many purposes, including inducing pathogen-related (PR) protein activity and attracting natural herbivore predators^67^. Treatment with (Z)-3-hexenol also increases *PR* gene expression, indicating that (Z)-3-hexenol directly promotes a series of defense responses. evk is released as a pest-induced plant volatile when *Pieris rapae* is fed on Arabidopsis^68^. Evk is a small, volatile chemical molecule with an α, β-unsaturated carbonyl structure. Due to its unique functional group, this molecule is dispersed rapidly and has a strong ability for long-distance diffusion, making it an effective signaling molecule. In the current study, for the first time, we found the key proteins that sense evk through molecular docking. The recognition of volatiles and receptors is the starting point of defense signals, which is the key to the transmission of defense signals between plants and the cause of population resistance. Based on the docking simulation results, it is predicted that evk will assist the electron transfer of RBOHD/RBOHF by binding to its FAD or NADPH binding sites (Figure 2). When the signal is recognized, we observed and characterized the whole process of evk mediated defense response: a. ROS burst; b. intracellular Ca^2+^ concentration change; c. WRKY53-CAM5 complex decomposing and release WRKY53; d. WRKY53 and TGA5 regulate expression of defense genes by evk.

Cytoplasmic free Ca^2+^ concentrations and H_2_O_2_ are thought to play crucial roles in the volatile-sensing mechanisms of plants, although these roles have not been directly demonstrated. We previously showed that evk is recognized by plants and induces H_2_O_2_ production, which ultimately mediates stomatal closure to reduce pathogen infection^69^. In the current study, we demonstrated that evk, a volatile compound released by plants, increased the levels of H_2_O_2_ and intracellular calcium in Arabidopsis mesophyll cells. CML8-ACA8 lowered the levels of intracellular calcium after the calcium ions have served their function, causing calcium levels in the cells to return to the resting state. H_2_O_2_ levels in mesophyll cells rose when Arabidopsis detected evk, with the H_2_O_2_ burst in leaves being mediated by NADPH oxidases. Therefore, H_2_O_2_ levels in mesophyll cells increased in an RBOHD- and RBOHF-dependent manner. Importantly, the evk-mediated ROS burst was unaffected by pretreatment with ruthenium red, which blocked membrane calcium fluxes (Figure 1a,1b).

Ca^2+^ homeostasis is tightly regulated in plant cells. The most abundant Ca^2+^ store in plant cells is the vacuole, with Ca^2+^ concentrations ranging from 0.2 to 5 mM^70^. Conversely, the chloroplast stroma contain <150 nM of free Ca^2+^, similar to that in the cytosol^71^. Evk treatment rapidly increased intracellular Ca^2+^ concentrations (Figure 3a,3b) and Ca^2+^ flow, but both effects were reduced in *rbohd rbohf* mesophyll cells (Figure 4a,4b). These results suggested that the H_2_O_2_ burst induced by evk took place prior to the rise in Ca^2+^ levels in Arabidopsis mesophyll cells. Additionally, the increase in intracellular calcium levels was less severe in the ruthenium red pretreatment group, indicating that the intracellular calcium pool plays a role in the increase in intracellular calcium concentrations mediated by evk. Similar observations were made following treatment with evk and acyclic compounds^72,73^. By contrast, (E)-2-hexenal, an electrophilic compound, mainly promotes Ca^2+^-influx from the extracellular space. Interestingly, ROS scavengers attenuated the (E)-2-hexenal-induced intracellular calcium transient, suggesting that ROS-dependent activation of plasma membrane Ca^2+^-permeable channels participates in the electrophile-dependent intracellular calcium transient^73^.

Ca^2+^ and H_2_O_2_ collaborate during two processes in plants: Ca^2+^-induced ROS generation and Ca^2+^-induced ROS release^29^. Here, we demonstrated how evk functioned as a signal transducer during ROS-induced Ca^2+^ release. Different stimuli are detected by various mechanisms throughout early signal transmission, making the sequence of early signal transmission events a unique “language” in plant resistance. Ca^2+^ is crucial for many signaling cascades, yet excessive amounts of Ca^2+^ in the cytoplasm are harmful to living organisms. The intracellular Ca^2+^ storage membrane and the plasma membrane both carry Type P ATPases, which take part in Ca^2+^ efflux. CaM can only be coupled with II B Ca^2+^-ATPases. CaM attaches to the N-terminal CaM-binding site when the N terminus of Ca^2+^ ATPases binds to Ca^2+74^. We discovered that mesophyll cells treated with evk had potent Ca^2+^ outflow by measuring the Ca^2+^ flux. We therefore investigated the possibility that the Ca^2+^ ATPase inhibitor eosin B would prevent this Ca^2+^efflux. Our findings demonstrated that Ca^2+^ ATPase contributed to the Ca^2+^efflux of mesophyll cells induced by evk and that Ca^2+^ efflux was blocked in *cml8* plants (Fig. 3c,3d). We hypothesize that the restoration of Ca^2+^ levels during the resting stage is mediated by an interaction between CML8 and ACA8 (Fig. 5a-5c).

Plants have developed sophisticated defense mechanisms against necrotrophic and biotrophic diseases^75^. Plants typically activate JA-induced defense responses against herbivorous insects or necrotrophic pathogens. Infection by *Pseudomonas syringae* pv. *tomato* DC3000 induces plants to accumulate SA, which promotes the interaction of NPR1 with TGA transcription factors and modulates the expression of pathogenesis-related (*PR*) genes. Plants activate SA-mediated resistance against biotrophic pathogens. *B. cinerea* infection activates JA biosynthesis and signaling pathways to promote the immune response^76^. These JA-related transcription factors then activate JA-responsive genes such as *PDF1*.*2* against necrotrophs^46^.

The cell redox status shifts due to the presence of the unsaturated carbonyl group of evk, which is highly oxidizable. The major plant hormone SA is extremely redox state sensitive, and evk triggers an H_2_O_2_ burst. Since there is a significant correlation between H_2_O_2_ and SA and WRKY53 is substantially activated by evk, as determined by RT-qPCR (Figure. 6d), we reasoned that evk might activate genes related to SA. We therefore performed RT-qPCR to examine the expression of all SA-related genes. evk treatment (particularly during the first 3 h) greatly stimulated the expression of SA-related genes (Figure 6a). This treatment also blocked the expression of several JA-related genes (Figure 6b). JA levels rapidly increased within 8 h of treatment, and SA accumulation increased dramatically at 3 h and remained unchanged at 8 h (Figure. 6c).

Interestingly, however, *PR1* was upregulated not only at 3 h but also at 8 h of evk treatment, suggesting that a regulator activates *PR* expression during both the early and late stages of treatment. As demonstrated by RT-qPCR, evk treatment of Arabidopsis leaves induced SA/JA antagonism, particularly the simultaneous activation of the defense genes *PR1* and *PDF1*.*2* at 8 h (Figure 6a,6b). evk treatment increased SA/JA levels in addition to triggering the antagonism of SA/JA. *PR1* and *PDF1*.*2* were previously found to be antagonistically activated; however, under evk treatment, they may become co-activated at a later time.

JA and SA act antagonistically to mediate defense responses: JA can repress SA-mediated defense^77^. SA also affects the expression of the genes encoding the JA biosynthesis enzymes LOX2 and AOS^78,79^. Several components of SA signaling also suppress JA signaling, including NPR1, the glutaredoxin GRX480, and class II TGA and WRKY transcription factors^79,80^. SA strongly reduces the accumulation of ORA59 (OCTADECANOID-RESPONSIVE ARABIDOPSIS AP2/ERF 59), a JA-responsive transcription factor, thereby inhibiting the expression of JA-responsive genes^80^. Biotrophic pathogen-induced SA accumulation inhibits CAT2 activity to inhibit JA accumulation by ACX2 and ACX3^81^.

In the current study, whereas SA-related gene expression was substantially elevated during the early stages of evk treatment, this expression was suppressed during the later stages, while JA-related genes were strongly induced. Whereas JA levels only increased during the late stage of evk treatment, SA was maintained at high levels, which were relatively constant in both the early and late stages. In particular, when *WRKY53* was mutated, we detected almost no *PR1* expression, whereas *PDF1*.*2* was significantly upregulated during the later stage of evk treatment (Figure. 7h,7i). Therefore, it is highly likely that WRKY53 serves as a node of SA/JA antagonism.

When *CAM5* or *WRKY53* was mutated, SA-related genes were downregulated (Figure. 7e-7g), indicating that CAM5-WRKY53 positively regulate the expression of these genes. In addition, CAM5 positively regulated *WRKY53*. These findings confirm the notion that CAM5 and WRKY53 interact and that the increase in calcium concentration causes WRKY53 to be released from the CAM5-WRKY53 complex (Figure. 8a-8d).

*PR1* was not significantly upregulated in *wrky53*, but *PDF1*.*2* was highly upregulated in this mutant. These findings indicate that WRKY53 is a key factor that positively regulates *PR1* expression and negatively regulates *PDF1*.*2* expression (Figure. 7h,7i). The dual-luciferase reporter assay further revealed that WRKY53 positively regulates *PR1* and negatively regulates *PDF1*.*2* transcription (Figure. 11c-11f). In an RT-qPCR assay, after evk treatment, the expression pattern of *WRKY53* showed a single peak, indicating that this gene was downregulated during the later stage of treatment, suggesting that other factors are involved in *PR* activation. TGA5 positively activated *PR1* expression. The WRKY53-CAM5 complex was unlocked when intracellular calcium levels rose following evk treatment (Figure. 8b). The freed WRKY53 was then able to bind to the *PR1* and *PDF1*.*2* promoters, increasing and inhibiting their transcriptional output, respectively (Figure 10a,10c; Figure 11c-11f). Additionally, the released WKY53 may influence the capacity of TGA5 to bind to *PR1* and *PDF1*.*2*. The capacity of TGA5 to bind to the as-1 motif in the *PR* promoter was enhanced by WRKY53 (Figure. 10b), whereas the ability of TGA5 to bind to this element in the *PDF1*.*2* promoter was inhibited by WRKY53 (Figure. 10d). As a result, WRKY53 greatly induced *PR1* expression while suppressing *PDF1*.*2* transcription. This study established for the first time that WRKY53 positively activates *PR1* transcription while negatively activating that of *PDF1*.*2*. However, WRKY53 reduced the binding capacity of TGA5 to the as-1 sequence and prevented the transcription of *PDF1*.*2* in the presence of both WRKY53 and TGA5. We discovered that *WRKY53* was expressed in a unimodal pattern, which explains why *PDF1*.*2* expression was suppressed during the early stages of evk treatment and was triggered when *WRKY53* expression decreased. *PR1* was undoubtedly activated during the early stages of evk treatment, and why it was still upregulated during the later stages of treatment, when *WRKY53* expression was lower, TGA5 still allows positive regulation of PR1 expression. Our findings support the hypothesis that a novel factor, TGA5, is involved in *PR1* activation and that TGA5 continues to positively regulate *PR1* expression during the later stage of treatment with evk.

Stress-related lipid oxygenation, whether enzymatic or non-enzymatic, will produce compounds with multiple molecular weights. However, the common feature of many of these compounds is the existence of highly conserved carbonyl groups in the molecule. There is a large subgroup of lipid oxidation products including α, β-unsaturated carbonyl. This kind of substance is also called reactive electrophile species (RES). Naturally occurring in plants α, β-unsaturated carbonyl compounds: evk, is an active small molecule released by plants under stress^19^. Through research, this paper proposes a framework that evk, as a RES substance, can achieve plant’s ‘REScue’ through complete defense response. The complete defense pathway is as follows: (1) recognition of evk (recognized by plant membrane protein RBOH); (2) Signal transduction (early signal generation: ROS burst, intracellular calcium concentration rise and intracellular calcium concentration recovery); (3) Expression of defense genes (positive regulation of PR1 gene by WRKY53 and TGA5, up-regulation of PDF1.2 gene); (4) accumulation of active substances (accumulation of hormone JA and SA in plants). When plants are faced with various kinds of stress, they will release a lot of volatiles. Little is known about the study of volatiles. This paper provides a valuable research idea for the study of other volatiles.

Evk-mediated ‘REScue’ helps to promote the survival of plants. In the early stage of defense, it can achieve the resistance to biotrophic pathogens represented by SA, and in the late stage of defense, it can achieve the resistance to herbivorous insects or necrotrophic pathogens by JA, thus improving the resistance of plants and promoting plant survival. In the future, it can be used as a potential green anti-insect and anti-bacterial drug released by pure natural plants to replace highly toxic synthetic drugs.

## Acknowledgments

This research was supported by the National Natural Science Foundation of China (31270655). We thank Meiqin Liu (Analysis and Testing Center, Beijing Forestry University) for her guidance on laser scanning confocal microscopy; Prof. Daoxin Xie (Tsinghua-Peking Joint Center for Life Sciences and MOE Key Laboratory of Bioinformatics, School of Life Sciences, Tsinghua University) for providing the plasmids: Cluc, Nluc pGreenII 0800-LUC and pGreenII 62-SK. All authors have approved the manuscript for publication, and no conflicts of interest exist.

## Author Contributions

Design of research: J.G.and Y.S. Performed experiments: J.G., Z.G., Z.W. and L.Y. Analysed data: J.G., S. M., C.Z., S.L.Wrote the manuscript: J.G. All authors have read and agreed to the published version of the manuscript. J.G., Z.G. and Z.W. contributed equally to this work.

## Data Availability Statement

The data that support the findings of this study are available from the corresponding author upon request.

## Author Contributions

Design of research: J.G.and Y.S.Performed experiments: J.G., Z. G. and Z. W. L.Y. Analysed data: J.G., S. M., C.Z., S.L.Wrote the manuscript: J.G. All authors have read and agreed to the published version of the manuscript. J.G., Z.G. and Z.W. contributed equally to this work.

