## Supplementary material for "CAM5, WRKY53, and TGA5 regulate defense gene expression mediated by the volatile organic compound ethyl vinyl ketone": supplymentary figure1-3

| Gene Name | Forward Primer Sequence (5'–3') | Reverse Primer Sequence (5'–3') |
| --- | --- | --- |
| *LOX2* | CACCTGACGAAGAGTACATTGG | GTAACACCATGCTCAGAGGTAGG |
| *PDF1.2* | CACCTGACGAAGAGTACATTGG | GTAACACCATGCTCAGAGGTAGG |
| *JMT* | CTAGGCAGAAGAGTAATGGAC | GTGAAGGCTCCGGCGAGG |
| *OPR1* | TTTACCCCTCCAAGACGGC | GAATCTCAACTCCATCAAAACCTG |
| *OPR3* | AAGGCAAGGGAGTGATGAGG | GAAAAAGGAGCCAAGAAAGGAT |
| *AOC3* | GAAGGAGATAGAAACAGTCCAGCA | CGAATCTGTCACCGCTCTTTT |
| *RBOHD* | ATCAAGGTGGCTGTTTACCC | GGGAGCTGATGTGATTGAGA |
| *PR1* | GGGGAAAACTTAGCCTGGG | CCACCATTGTTACACCTCACTTT |
| *NPR1* | GCTTGCGGAGAAGACGACA | CCGACGACGATGAGAGAGTTTA |
| *ICS1* | CTGGGCTCAAACACTAAAACACA | GCGGAGATTAGCCTTTCAGTG |
| *NPR4* | GCATTGAGAGAGTGGCGAGA | GCTTCACGAGTTCTACATCATCTG |
| *PAD4* | TTCAGTTGGGGAATCTTGTGG | AAAGTGCGGTGAAAGCGG |
| *SARD* | CGTAAGTTTAGAATCGGTGCG | TTGATGTGGCGAGAGGAGAG |
| *WRKY53* | GACGGCTGTTGCTGAGACTAA | CAAACTCTTCACTTCTCGGACTTC |
| *CAM5* | ATGGCAGATCAGCTCACCGA | GAGAATACGGCAGTGACTTTTCC |
| *ACTIN2* | AGTGGTCGTACAACCGGTATTGT | GATGGCATGAGGAAGAGAGAAAC |
| *EF1*α | TCCAGCTAAGGGTGCC | GGTGGGTACTCGGAGA |

**Supplementary Figure 1.**

**Supplementary Figure 2.**

| w-boxes | Probe sequence (5'*-*3') |
| --- | --- |
| *PDF1*.*2* as-1-F | ATGTTGTATTTGTTCGACGATGACGAAGGT |
| *PDF1*.*2* as-1-R | ACCTTCGTCATCGTCGAACAAATACAACAT |
| *PDF1*.*2* w-box-F | ATAATTATTCTTGACTGATGTATGCATATA |
| *PDF1*.*2* w-box-R | TATATGCATACATCAGTCAAGAATAATTAT |
| *PR1* w-box-F | TTTATTTGAAAATTGACTGTAGATATAAAC |
| *PR1* w-box-R | GTTTATATCTACAGTCAATTTTCAAATAAA |

**Supplementary Figure 3.**

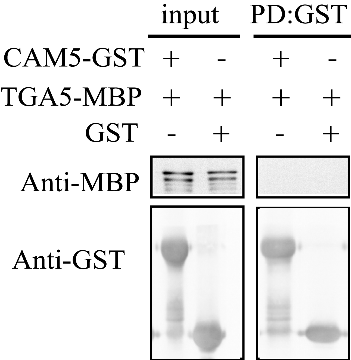

CAM5 do not interact with TGA5. Since WRKY53 can interact with TGA5, and when intracellular calcium ions are activated and raised by evk and WRKY53-CAM5 complex is untied, WRKY53 and TGA5 can interact. So whether TGA5 and CAM5 can also interact? We conducted Pull Down experiments and found that CAM5-His can not be pulled down by TGA5-GST.
